# Low land-use intensity buffers grasslands against future climate and inter-annual climate variability in a large field experiment

**DOI:** 10.1101/2023.12.20.572519

**Authors:** Lotte Korell, Martin Andrzejak, Sigrid Berger, Walter Durka, Sylvia Haider, Isabell Hensen, Yva Herion, Johannes Höfner, Liana Kindermann, Stefan Klotz, Tiffany M. Knight, Anja Linstädter, Anna-Maria Madaj, Ines Merbach, Stefan Michalski, Carolin Plos, Christiane Roscher, Martin Schädler, Erik Welk, Harald Auge

## Abstract

Climate and land-use change are key drivers of global change. Full-factorial field experiments in which both drivers are manipulated are essential to understand and predict their potentially interactive effects on the structure and functioning of grassland ecosystems. Here, we present eight years of data on grassland dynamics from the Global Change Experimental Facility (GCEF) in Central Germany. On large experimental plots, temperature and seasonal patterns of precipitation are manipulated by superimposing regional climate model projections onto background climate variability. Climate manipulation is factorially crossed with agricultural land-use scenarios, including intensively used meadows and extensively used (i.e. low-intensity) meadows and pastures. Inter-annual variation of background climate during our study years was high, including three of the driest years on record for our region. The effects of this temporal variability far exceeded the effects of the experimentally imposed climate change on plant species diversity and productivity, especially in the intensively used, species-poor grasslands. These changes in productivity and diversity in response to alterations in climate were due to immigrant species replacing the target forage cultivars. This shift from forage cultivars to immigrant species may impose additional economic costs in terms of a decreasing fodder value and the need for more frequent management measures. In contrast, the extensively used, species-rich grasslands showed weaker responses to both experimentally manipulated future climate and inter-annual climate variability, suggesting that these diverse grasslands are more resistant to climate change than intensively used, species-poor grasslands. We therefore conclude that an extensive management of agricultural grasslands, together with other measures to increase species diversity, can stabilize primary productivity under climate change.

## Introduction

Grasslands cover more than one third of the Earth’s terrestrial surface and provide important ecosystem functions and services (Bengtsson et al. 2019). They store at least one third of the terrestrial carbon (White et al. 2000, Bai and Cotrufo 2022), are essential for food production (O’Mara 2012), and can maintain an exceptionally high biodiversity at a small spatial scale (Biurrun et al. 2021, Sabatini et al. 2022). Climate change and land-use intensification - two main constituents of anthropogenic global change – are major threats for the biodiversity and functioning of these important ecosystems (Gibson and Newman 2019, Schils et al. 2022).

Over the last decades, global mean temperature has increased by about 1°C, and is projected to increase by 1.5 to 4°C (2070-2100) depending on the emissions scenario (IPCC 2021). In addition, seasonal precipitation patterns (e.g. intra- and inter-annual variability) have changed across the globe, a trend which is expected to continue in the future (IPCC 2021). As a consequence, extreme climate events, such as droughts, have dramatically increased in frequency and extent over the last decades, and are expected to do so even more in the future (IPCC 2021). Climate change experiments that monitor community responses can establish cause-effect relationships and can improve our mechanistic understanding of the consequences of anthropogenic climate change on community- and ecosystem-level responses.

Climate change can lead to changes in the phenology, demography and abundance of plant species (Piao et al. 2019, Morris et al. 2020, Compagnoni et al. 2021), subsequently leading to alterations in interactions within and across trophic levels (Gornish and Tylianakis 2013, Renner and Zohner 2018), ultimately driving shifts in plant community attributes (Parmesan 2006, Bellard et al. 2012). Results from climate warming experiments show that warming can alter plant community composition and cause declines in species richness (Gruner et al. 2017, Bastazini et al. 2021). Conflicting results exist on the effect of warming on annual aboveground productivity (see meta-analysis by Wu et al. 2011, Song et al. 2019). Meta-analysis show that experimental decreases in mean annual precipitation usually cause declines in biodiversity and productivity, whereas experimental increases in mean annual precipitation lead to higher biodiversity and productivity in terrestrial plant communities (Wu et al. 2011, Song et al. 2019, Korell et al. 2021). Changes in the (seasonal) pattern of precipitation, which do not necessarily correspond with large changes in the mean, are still understudied (Jentsch et al. 2007, Korell et al. 2020, Hajek and Knapp 2022). Yet, even small changes in seasonal precipitation patterns can have profound impacts on ecosystem processes such as productivity (Fay et al. 2003, Wilcox et al. 2015, Gherardi and Sala 2019) and can lead to changes in the composition and diversity of plant communities (Knapp et al. 2020). Despite anthropogenic climate change comprises simultaneous alterations in temperature and precipitation, their combined effects have been rarely considered in experimental studies so far (Wu et al. 2011, Song et al. 2019). A predictive understanding of plant community and ecosystem responses requires experiments that cover future climate scenarios (Korell et al. 2020), e.g. manipulating changes in seasonal precipitation patterns in combination with alterations in temperature (Schädler et al. 2019).

Climate change is acting in concert with other components of global change (Sala et al. 2000, Komatsu et al. 2019, Speißer et al. 2022) such as land-use change. Semi-natural grasslands in Central Europe have historically been created by human activities and therefore need regular management practices, i.e. mowing and/or grazing, to be maintained (Poschlod et al. 2009). At low management intensity (= extensively used) grasslands are usually highly diverse (Tälle et al. 2016). Yet, many European grasslands have been converted to intensively used grasslands through high fertilizer input, higher mowing frequency and/or livestock density, and sowing of highly productive grass cultivars (Poschlod et al. 2005). This land-use intensification has led to a conversion of less productive and species-rich, extensively used grasslands to highly productive but species-poor, intensively used grasslands (Gossner et al. 2016, Beckmann et al. 2019). According to the insurance hypothesis in biodiversity research, more diverse ecosystems are suggested to show higher resistance (i.e., less change in ecosystem responses, Pimm 1984) against perturbations compared to species-poor ecosystems (Yachi and Loreau 1999). The few experimental studies available indicate that species-poor, intensively used grasslands are less resistant to drought events compared to less productive, species rich, extensively used meadows (Vogel et al. 2012, Stampfli et al. 2018, Isbell et al. 2021), but this has not yet been tested using realistic scenarios of land use and future climate.

Community and ecosystem responses to environmental change are known to depend on the temporal scale (Smith et al. 2009). Initially, changes have large effects on resident communities that evolved and assembled under previously existing environments. These effects dissipate over time until the community arrives at a new state (Hastings 2004, Sandel et al. 2010). According to the hierarchical response framework (HRF, Smith et al. 2009), such transient responses should be characterized by a temporal hierarchy of mechanisms beginning with responses at the level of individuals, followed by species reordering, and finally species loss and immigration. Immigration is expected to lead to particularly large changes in ecosystem properties such as aboveground productivity (Smith et al. 2009, Alexander et al. 2015). However, land-use intensity may affect how many species can potentially immigrate into grasslands systems, e.g. due to differences in management practices, fertilizer input, and sowing of species-poor seed mixtures, ultimately mediating the response of such grasslands to climate change.

Here, we present the results of eight years (2015-2022) of realistic climate manipulations on the productivity, diversity and composition of grasslands of different land-use intensity. We used the Global Change Experimental Facility (GCEF), a large experimental platform located in Central Germany. Future temperature and seasonal precipitation are experimentally manipulated based on regional climate model projections and factorially crossed with different realistic agricultural land-use scenarios, including intensively used meadows and extensively used meadows and pastures (Schädler et al. 2019). As background climate conditions varied considerably during our study period, including three years of extreme drought (Rakovec et al. 2022), we could also compare community responses to increases in both experimental and background inter-annual climate variability. Such high climate variability can be also considered a consequence of anthropogenic global change (Naumann et al., 2021).

Specifically, we asked the following questions: (1) How does future climate affect plant species diversity of differently managed grasslands? We expect that highly productive cultivars in intensively used grasslands may be less tolerant to summer droughts than resident species in extensively used grasslands. A resulting greater dieback of these cultivar species in intensively used grasslands may allow more species from the regional species pool to immigrate, especially if these immigrating species are better adapted to future climate conditions, resulting in a net increase in species richness under future climate conditions. (2) How does future climate affect productivity of differently managed grasslands? Climate manipulation may lead to an overall decline in productivity and responses will be stronger in the intensively used grasslands due to less drought tolerant cultivars. (3) What is the relative importance of inter-annual background climate variability and experimental climate change on productivity, and how do grasslands differ in their resistance against background climate variability? We hypothesize that inter-annual climate variability, including extreme drought years, will have relatively more drastic effects on productivity compared to effects of the climate manipulation, which includes wetter springs and falls that could counteract negative effects of drier summer conditions. We expect intensively used grasslands to show a lower resistance to inter-annual climate variability than extensively used ones.

## Materials and Methods

### Study site and experimental design

The experimental site is located at the Field Experimental Station of the Helmholtz-Centre for Environmental Research – UFZ in Bad Lauchstädt, Central Germany (51°22’60 N, 11°50’60 E, 118 m a.s.l.). It is characterized by a sub-continental climate with a mean annual temperature of 8.9°C and a mean annual precipitation of 489 mm (1896-2013). During the study period (2015-2022), which included three drought years in a row (2018-2020), annual precipitation was on average 97 mm lower, and annual temperature was 1.8 K warmer than the long-term mean (supporting information, fig. S1). The nutrient-rich soil is a Haplic Chernozem upon carbonatic loess substrate, and has a high humus content as well as a high water holding capacity (for further details see Altermann et al. 2005).

The Global Change Experimental Facility (GCEF) was established on a former arable field in 2012. The experiment was set up using a two-factorial split-plot design: 10 main plots of 80 m x 24 m size, which have a minimum distance of 25 m to each other, were randomly assigned to one of two climate treatments (ambient climate vs. future climate). Each main plot was subdivided into five subplots of 16 m x 24 m, which in turn were randomly assigned to one of five land-use treatments (organic farming, conventional farming, intensively used meadow, extensively used meadow, extensively used pasture). This results in a total of 50 subplots with five replicates for each combination of climate and land use. In 2013, oat was sown on all plots in order to homogenize soil conditions including soil seed bank. In this study, we will consider the three grassland land uses only: the intensively used meadows (IM) were set up in fall 2013, and the extensively used meadows (EM) as well as the extensively used pastures (EP) were sown in in early spring 2014. Climate manipulation started in spring 2014 for all land-use treatments (see Schädler et al. 2019 for details).

### Climate manipulation

Within the GCEF, climate was manipulated according to projections of dynamic regional climate models for the period of 2070–2100. We used the mean projection of 12 simulations based on three models and four emission scenarios, which corresponds to an increase in temperature by 2 K and a changed seasonal pattern of precipitation, with about 10% increase in spring and fall and about 20% decrease in summer (see Schädler et al. 2019 for details). We superimposed the seasonal change in precipitation on the ambient precipitation pattern, thereby retaining the background climate variability. Roof structures were installed above each main plot representing future climate, equipped with translucent tarpaulins which closed every day from sunset to sunrise to achieve passive warming. We considered passive night-time warming to be in accordance with the predicted asymmetry in global warming between day and night (Harvey 1995, Davy et al. 2017). However, the resulting temperature increase was lower than projected: e.g., mean daily temperature increased by 0.55 K at a height of 5 cm and by 0.24 K at a height of 70 cm (see Schädler et al. 2019 for details). In the summer months (June to August), roofs were additionally closed during daytime rain events to reduce summer precipitation by 20%. In spring (March to May) and fall (September to November) stored rainwater was added using an irrigation system to increase precipitation by 10%. Air and soil temperature as well as precipitation are continuously measured on the 10 main plots representing ambient and future climate treatments (Schädler et al. 2019). Plots of ambient climate are equipped with similar roof constructions, but without tarpaulins and irrigation system, to mimic possible side effects (Kreyling et al. 2017).

### Land-use treatments

The three grassland types differ with respect to a range of factors, such as initial species composition, fertilization, management type, and management frequency. Considering the different management, we will refer to them as land-use types according to Schädler et al (2019) although they represent the same land-use cover (i.e. grassland). For the extensively used meadows (EM) and extensively used pastures (EP), seeds of 56 native grassland species (14 grasses, 10 legumes, 32 non-legume forbs; table S1) from the regional species pool were sampled in one to six local populations per species. The selected species are typical for mesophilous to dry grasslands as well as steppe grasslands of Central Germany (Arrhenatherion, Cynosurion, Festucion valesiacae, and Cirsio-Brachypodion communities according to Schubert et al. 2010). In early March 2014, seeds were sown at a total density of c. 1,600 seeds m^−2^ (2.2 g m^−2^), which is within the range recommended for grassland restoration in the region (Kirmer and Tischew 2006). A few poorly germinated forb species were re-sown in early October 2014. In summer 2014, the first growing season after sowing, mowing was applied once to EM and EP in order to suppress weed establishment.

Beginning in 2015, the year when vegetation sampling started, EM was mown twice every year (late spring, midsummer), except in 2018, 2019 and 2020, when mowing had to be suspended in summer due to insufficient regrowth. EP was mown in late spring 2015 as well, and first grazing by sheep was initiated in midsummer 2015. The lower grazing intensity at the beginning of the experiment was intended to facilitate establishment of sown species. In 2016, grazing took place two times, in late spring and midsummer. Starting in 2017, grazing frequency was increased to three times per year (early spring, late spring, midsummer). In 2018, 2019, and 2020, however, grazing had to be reduced to two times each year, and in 2022 to one time, because of insufficient regrowth owing to summer drought. Grazing is conducted as short-time, high-intensity grazing with a group of 10 to 13 adult sheep and 10 to 20 lambs remaining on each EP subplot for 24 h. Such grazing management is recommended for nature conservation purposes (Landesamt für Landwirtschaft 2010).

For the intensively used meadows (IM), species composition follows official recommendations of state authorities for drier sites (Landesanstalt für Landwirtschaft und Gartenbau Sachsen-Anhalt 2020). The recommended seed mixture “RG8” contains 5 grass cultivars: 10% of early-season *Lolium perenne* cv. ‘Nui’, 10% of midseason *L. perenne* cv. ‘Toronto’, 50% of ‘Festulolium’ (*Festuca pratensis* x *Lolium multiflorum*) cv. ‘Felopa’, 20% of *Dactylis glomerata* cv. ‘Lidacta’, and 10% of *Poa pratensis* cv. ‘Balin’, with a total sowing density of 3.0 g m^−2^. The IM grassland was sown in fall 2013 and mown three times in 2014 (midspring, early summer, early fall). Starting in 2015, mowing frequency was increased to four times each year (midspring, late spring, midsummer, midfall). Owing to dry summers, mowing frequency had to be reduced to 2 cuts each year in 2018, 2019, and 2020, and to three cuts in 2022. In fall 2020, IM subplots were plowed and re-sown using the same seed mixture as before. This is part of the typical management of intensively used grasslands in Central Europe to re-establish the desired mixture of cultivars. According to agricultural practice, mineral fertilizer is applied every year at the start of the growing season (90 kg/ha N, 30 kg/ha P, 110 kg/ha K) and after the first (80 kg/ha N), second (70 kg/ha N), and third cut (60 kg/ha N).

### Vegetation sampling and compilation of derived variables

A 3 m x 3 m permanent quadrat was established on each of the 30 grassland subplots. On this quadrat, percentage cover of each higher plant species was visually estimated two times each year on EM and EP (late spring and midsummer, before mowing or grazing) and four times each year on IM (midspring, late spring, midsummer, midfall), except in 2015 and 2020 when IM was sampled three times per year only. Data on species cover were used to calculate total species richness, richness of sown species (hereafter referred to as “resident species”), richness of not-sown species (hereafter referred to as “immigrant species”), as well as average cover of each species per 9 m^2^ sampling quadrat and year. As a measure of community evenness we used a bias-corrected version (Chase et al. 2018) of the probability of interspecific encounter, PIE (Hurlbert 1971). We used the average cover of species per year rather than the number of individuals and converted PIE into the effective number of species 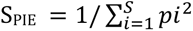, where S is the total number of species and *pi* the proportion of each species *i* and it is equivalent to 1/Simpson’s index (Hill 1973, Jost 2006).

To measure standing biomass of each plant species, aboveground biomass samples were taken just after cover estimation, i.e. two times a year on EM and EP, and four times a year on IM. Aboveground biomass was cut with scissors at a height of about 2 cm within four 20 cm x 50 cm metal frames which were randomly placed in two 3 m x 3 m areas next to the area used for cover estimates within each grassland subplot. Biomass samples were then sorted by species and dead biomass, dried at 70 °C for 48 h, and weighed. Only three biomass harvests took place on IM subplots in 2018 because there was no regrowth prior to the fourth occasion, and in 2020 because plots were plowed and re-seeded. Species-specific dry masses of the four samples per subplot were averaged to one biomass value per species, harvest date and subplot, and then used to calculate average biomass of resident and immigrant species per year for each subplot.

To estimate standing biomass and annual above-ground net primary production (ANPP), we had to apply different methods to EM and IM, and to EP, respectively (see Ruppert and Linstädter 2014 for method comparison). For EM and IM subplots, we used the biomass data described above and calculated standing biomass for each harvest date by adding species-specific biomass and dead biomass. As EM and IM were completely mown on each occasion, standing above-ground biomass of subsequent harvest dates was considered to represent regrowth. To estimate ANPP, we therefore added standing biomass of subsequent harvest dates of a given year. For EP subplots, additional biomass samples were taken in parallel to each grazing event using “moveable exclosures” (McNaughton et al. 1996): Four cages of c. 1 m² area were randomly placed on each EP subplot to exclude vegetation underneath from grazing. Immediately after grazing, cages were removed and biomass was harvested within 20 cm x 50 cm metal frames as described above. Similar biomass samples were taken at four randomly chosen positions outside cages on each subplot. Samples were dried at 70 °C for 48 h, and weighed. Biomass values of the four samples inside cages and of the four samples outside cages were averaged for each EP subplot to estimate standing biomass on each grazing event and remaining biomass after grazing, respectively. Re-growth between two grazing events was calculated as the difference in (grazed) biomass outside cages at time t and (ungrazed) biomass inside cages at time t+1. ANPP was then calculated by adding standing biomass measured on the first grazing date and regrowth during the following grazing events in each year.

To characterize climatic conditions, we measured precipitation and air temperature at a height of 1.5 m and 70 cm above soil surface, respectively, on each main plot. We compiled precipitation sum and mean air temperature for each month, for the four seasons spring (March to May), summer (June to August), fall (September to November) and winter (December to February), and for each study year. Because precipitation and temperature were correlated (r = −0.90; P < 0.01), we used De Martonne’s index dMI which combines the two measures (Andrade et al. 2021) to characterize drought conditions for ambient and future climate treatments on an annual basis: dMI = precipitation [mm] / (10 + temperature [°C]).

### Statistical analyses

To analyze plant species diversity and productivity, generalized linear mixed effects models were fitted to the data using the procedure GLIMMIX in SAS v. 9.4 (© SAS Institute Inc., Cary, NC, USA; see supporting information B for SAS Code). For count data (total species richness, richness of residents and richness of immigrants) we used models with Poisson distribution and log link function; for all biomass values (annual productivity, standing biomass, biomass of residents and biomass of immigrants) models with log-normal distribution and identity link function; and for proportional richness of immigrant species (richness of immigrants / total species richness) models with binomial distribution and logit link function. As proportional biomass of immigrants (biomass of immigrants / total living biomass) was not based on count data, we applied a logit-transformation to the data and used models with Gaussian distribution (Warton and Hui 2011). We empirically applied a log transformation to S_PIE_ values, and used a model with Gaussian distribution as well.

As we were interested in community dynamics across the experimental period, we considered the eight study years explicitly. Since measurements taken on the same experimental units across time are not independent, we applied repeated measurement analyses which differentiate between within-subject effects and between-subject effects (von Ende 2001). According to the split-plot design of the experiment, the between-subject model included the fixed factor climate at the main-plot level and the random effect of main plot nested within climate as the respective error term, while the factor land use and the climate x land use interaction were considered as fixed effects at the subplot level. The within-subject model included the fixed effects of year and of the year x climate interaction at the main-plot level, and the random effect of year x main plot nested within climate as the respective error term, as well as the year x land use and year x climate x land use interaction as fixed effects at the subplot level. We included a first-order autoregressive covariance structure, if the 95% confidence intervals showed a significant temporal autocorrelation or if the Akaike Information Criterion revealed an improved model fit. If appropriate, we explicitly accounted for heteroscedasticity among land-use types, and for over- or underdispersion by including a group-specific multiplicative dispersion parameter. In order to test a-priori hypotheses of differences between the intensively used grassland and the two extensively used grasslands, we decomposed the land use effect into orthogonal contrasts. In order to test how land-use types differed in their response to climate manipulation, we “sliced” the climate x land use interaction into the simple main effect (Woodward and Bonett 1991) of climate on each level of land use. For graphical presentation, we extracted the marginal means and their standard errors from the mixed-effect models.

In order to investigate how productivity of the three land-use types depended on inter-annual variability in climatic conditions, and how experimental climate manipulation affected this relationship, we applied an ANCOVA to mean ANPP, with treatment-specific annual dMI, land use, climate and their interactions as fixed effects; year was included as random effect in order to take into account that measurements taken in one year are not independent. To quantify the resulting temporal stability in productivity, we calculated the coefficient of variation (CV) of ANPP for each of the 30 climate x land use subplots across the eight study years. We are aware of the disadvantages of CV as an inverse measure of stability (Reckling et al. 2021), however the time frame of eight years was too short to allow for the commonly recommended corrections (e.g. Döring and Reckling 2018). We empirically applied a square root transformation to CV, and used a model with Gaussian distribution. According to the split-plot experimental design, climate was considered as fixed effect and main plot nested within climate as random effect at the main-plot level, and land use as well as the climate x land use interaction as fixed effects at the subplot level. To test our hypothesis that the extensively used grasslands are less variable in time (i.e. more stable) than intensively used grasslands, we decomposed the climate x land use interaction into the respective contrasts.

To compare response of ANPP to experimentally imposed climate change with response of ANPP to inter-annual climate fluctuations, we first calculated the log response ratios (LRR) (Hedges et al. 1999) of annual aboveground productivity between each subplot under future climate and the closest subplot under ambient climate for each land-use type and each year. The LRR values per subplot were averaged across years, and the mean among the five replicates and its standard error were calculated for each land-use type. We then compiled the LRR between the driest year (2018) and the wettest year (2021) for each subplot of the ambient climate treatment, and compared the mean response of each land-use type with its response to the experimental climate manipulation (see Kröel-Dulay et al. 2022). Annual means of the land-use specific LRR between future and ambient climate were furthermore used to test if magnitude and direction of response to experimental climate manipulation depend on background climatic conditions of each year, using an ANCOVA with annual dMI, land use and their interaction as fixed effects and year as random effect.

## Results

### Magnitudes of climate manipulations and of inter-annual climate fluctuations

Experimentally imposed climate change resulted on average in a 6.9 ± 0.7 mm increase (+11.0%) of precipitation during spring (March to May), in a 25.5 ± 4.6 mm reduction (−21.4%) during summer (June to August), a 5.5 ± 0.6 mm increase (+6.0%) during fall (September to November), and unintentionally in a 4.8 ± 1.0 mm decrease (−7.4%) in winter (December to February) relative to plots of ambient climatic conditions (means ± standard error). Consequently, annual precipitation was reduced by on average 18.2 ± 4.0 mm (−5.2%) under future climatic conditions (fig. 1). Experimental climate change increased air temperature by on average 0.14 ± 0.01 K in spring, 0.21 ± 0.01 K in summer, 0.21 ± 0.01 K in fall, and 0.08 ± 0.00 K in winter, resulting in a mean annual increase of 0.16 ± 0.01 K. During the study period, annual ambient precipitation (328 ± 26 mm year^−1^) was on average 161 mm lower than the long-term mean of 489 mm. However, it varied considerably among study years (range: 209 mm to 460 mm) with 2018, 2019, 2020, and 2022 being exceptionally dry years (48%, 28%, 22% and 27% less than the long-term mean, fig. 1). Mean annual temperature (10.8 ± 0.2 °C, range: 9.8 °C to 11.3 °C) was on average 1.9 K warmer than the long-term mean of 8.9 °C.

**Fig. 1:**
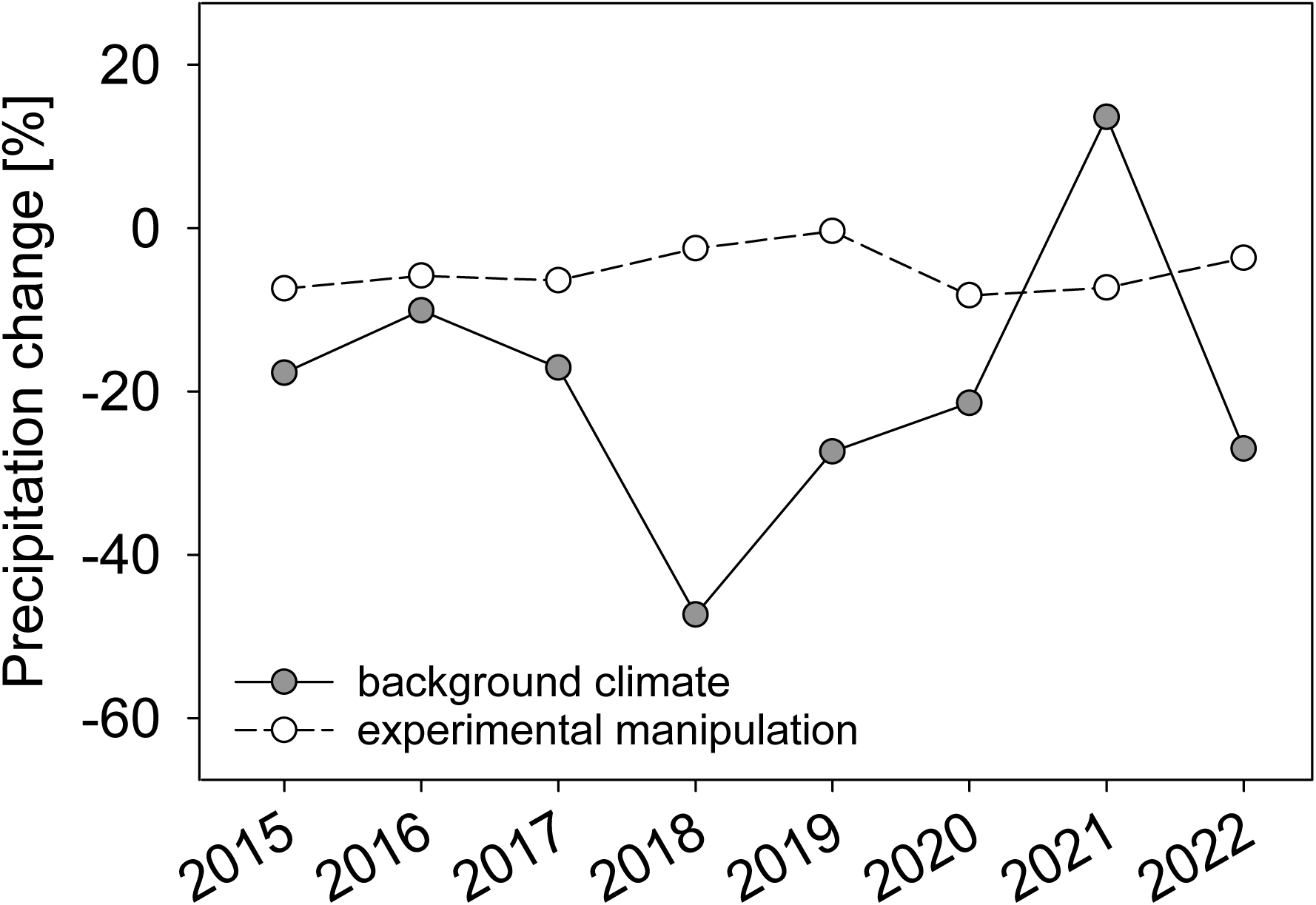
Effect of climate manipulation on annual precipitation (relative differences between future and ambient climate treatments), and inter-annual variation in background precipitation (relative differences between annual precipitation and the long-term mean 1896-2013) across the eight study years. Please note that the mean effect of climate manipulation does not reflect the seasonal change in precipitation pattern (increase in spring and fall, decrease in summer).

### Effects of experimental climate change on plant diversity across study years

Intensively used meadows (IM) showed, on average, a lower species richness (13.7 ± 0.3; mean ± standard error of 9 m^2^ sampling plots across years) than the two extensively used grasslands. Among them, species richness was higher in the extensively used meadows (EM; 46.4 ± 0.6) compared with the extensively used pastures (EP; 43.1 ± 0.8) (significant land use contrasts, table 1). The effect of experimental climate change on species richness differed among land-use types and among years (significant three-way interaction). While species richness of EM and EP showed only a minor, positive response to future climate during the first study years, richness of IM responded more strongly (fig. 2a). However, this response was not consistent across years: Richness of IM was elevated under future climatic conditions during the first study years, but reduced in 2020 before re-sowing. Generally, species richness of EM and EP showed rather small inter-annual variability, with a decrease during the drought years and a slight recovery afterwards. For IM, species richness increased substantially in the first years following establishment, declined in the first drought year 2018 but increased strongly in the second drought year 2019.

**Fig. 2:**
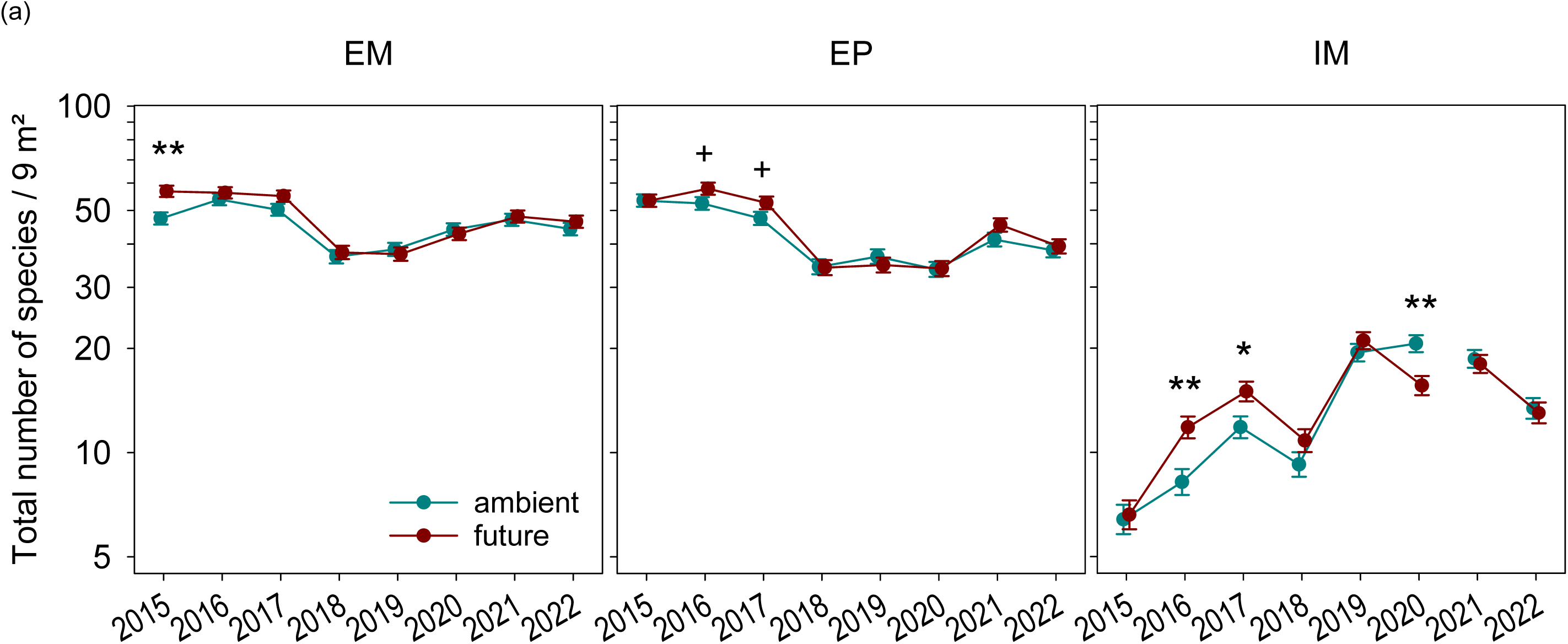
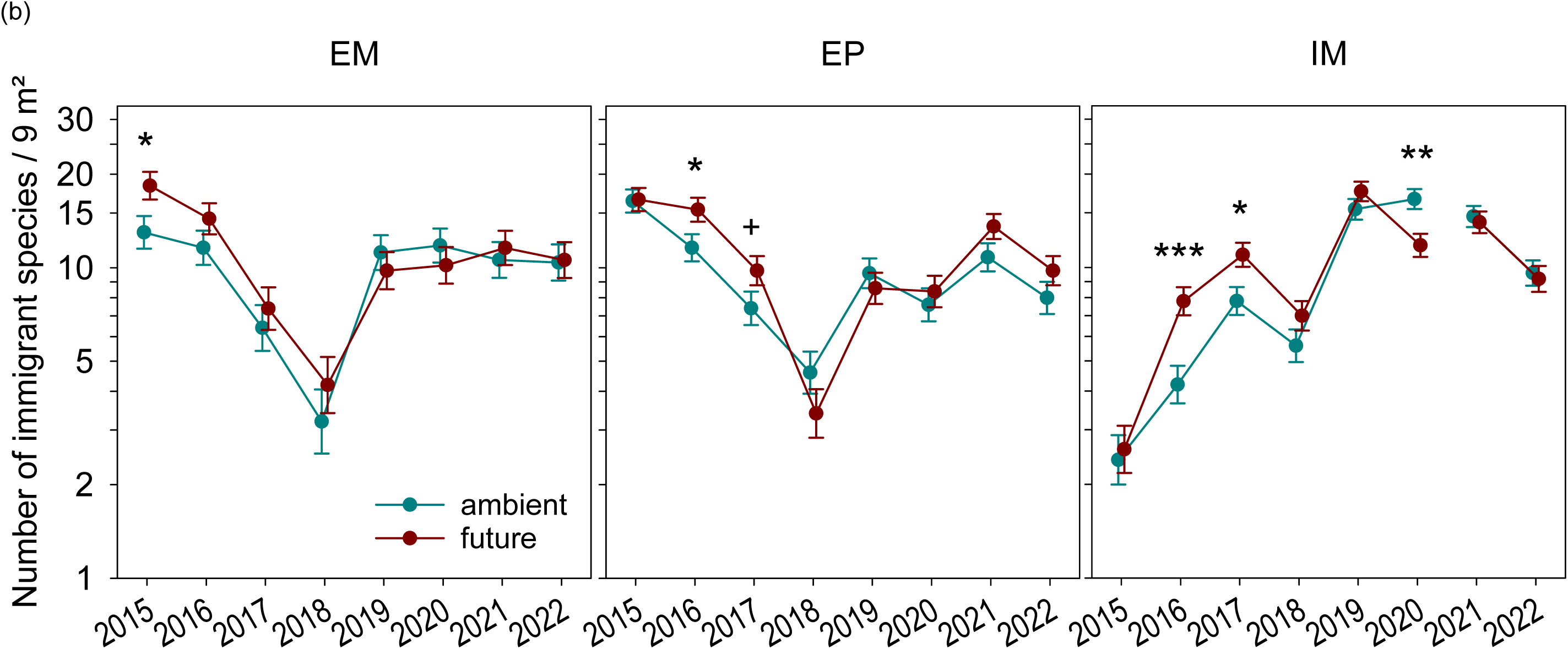
(a) Total plant species richness, and (b) species richness of immigrants of extensively used meadows (EM), extensively used pastures (EP) and intensively used meadows (IM) under ambient and future climatic conditions across eight study years, as measured on 9 m^2^ sampling plots (y axes log scaled). Please note that IM was re-sown in fall 2020. (Marginal means and their standard errors extracted from generalized linear mixed effects models; significance of simple main effects of climate for each year and land-use type: + p < 0.1, * p < 0.05, ** p < 0.01, *** p < 0.001.)

**Table 1:** Results of repeated measurements ANOVAs based on generalized linear mixed effects models for the effects of climate, land use and year on total plant species richness, species richness of immigrants, annual above-ground net primary production (ANPP), and proportional contribution of immigrant species to total above-ground biomass. To test a-priori hypotheses, the main effect of land use is decomposed into orthogonal contrasts, and the climate x land use interaction is decomposed into simple main effects of climate on each land use type (EM: extensively used meadows, EP: extensively used pastures, IM: intensively used meadows; d.f. enumerator and denominator degrees of freedom; + p < 0.1, * p < 0.05, ** p < 0.01, *** p < 0.001).

|  | d.f. | F ratio |  |  |  |
| --- | --- | --- | --- | --- | --- |
|  |  | Total richness | Richness of immigrants | ANPP | Proportional biomass of immigrants |
| Climate | 1, 8 | 4.12 <sup>+</sup> | 3.84 <sup>+</sup> | 3.56 <sup>+</sup> | 0.58 |
| Land use | 2, 16 | 1029.59 *** | 4.99 * | 705.80 *** | 5.05 * |
| Extensive vs. Intensive | 1, 16 | 2027.29 *** | 9.96 ** | 1411.26 *** | 0.16 |
| EM vs. EP | 1, 16 | 17.14 *** | 0.18 | 68.46 *** | 9.70 ** |
| Climate x Land use | 2, 16 | 0.16 | 0.10 | 1.45 | 4.54 * |
| Climate \| EM | 1, 16 | 2.41 | 1.49 | 0.13 | 0.94 |
| Climate \| EP | 1, 16 | 1.16 | 1.58 | 4.89 * | 0.08 |
| Climate \| IM | 1, 16 | 1.62 | 3.36 <sup>+</sup> | 3.41 <sup>+</sup> | 4.81 * |
| Year | 7, 56 | 28.25 *** | 27.69 *** | 27.87 *** | 10.14 *** |
| Year x Climate | 7, 56 | 2.82 * | 3.01 ** | 1.14 | 1.55 |
| Year x Land use | 14, 112 | 35.65 *** | 27.01 *** | 8.01 *** | 22.59 *** |
| Year x Climate x Land use | 14, 112 | 2.58 ** | 1.63 <sup>+</sup> | 0.92 | 1.57 <sup>+</sup> |

Consistent with differences in the numbers of species sown, realized richness of sown species (“residents”) was much lower on the intensively used grassland (3.9 ± 0.0) than on the two extensively used grasslands. Among the two extensively used grasslands, richness of residents was larger on EM (36.1 ± 0.4) compared to EP (33.0 ± 0.5; table S2). Furthermore, resident species richness showed larger temporal fluctuations on EM and EP compared to IM (significant year x land use interaction) but weak response to climate treatments (fig. S2a).

A considerable number of not-sown species recruited by immigration from the regional species pool or from the soil seed bank (“immigrants”). While species richness of immigrants was, on average, slightly lower on IM (9.8 ± 0.3) than on EM (10.3 ± 0.5) and EP (10.1 ± 0.5; table 1), it showed a strong increase on IM and a decline on EM and EP during the first study years (significant year x land use interaction; table 1, fig. 2b). After a general decline in the first year of drought, the number of immigrant species increased again during the following year in all three grassland types. In contrast to resident richness, richness of immigrants was higher under future climate during the first study years but this effect vanished in subsequent years (significant year x climate interaction; table 1, fig. 2b). The resulting proportional contribution of immigrants to total richness was much larger on IM than on EM and EP, but besides that, it mirrored their absolute numbers (table S2, fig. S2b).

The effective number of species, S_PIE_, as a measure of evenness was on average higher on the two extensively used grasslands compared to the intensively used grassland, but there were also striking differences between them in the temporal dynamics of S_PIE_ (significant year x land use interaction; table S2, fig. S2c). On EM and EP, S_PIE_ decreased during the first four to six study years, while on IM, a slight drop at the beginning was followed by a strong increase in S_PIE_ until plots were re-sown. The effect of experimental climate change on S_PIE_ depended on land use as well (significant climate x land use interaction). On EM, S_PIE_ was on average higher under future climatic conditions compared to ambient conditions, but did not significantly differ among climate treatments on EP and IM.

### Effects of experimental climate change on ANPP across study years

Annual aboveground net primary production (ANPP) of the intensively used grassland (1,015 ± 21 g m^−2^ year^−1^ across the study period) was about twice as large as productivity of the two extensively used grasslands, and was higher on EM (553 ± 18 g m^−2^ year^−1^) than on EP (405 ± 14 g m^−2^ year; significant land use contrasts, table 1). In addition, ANPP of IM showed a much steeper decline during the drought years and recovered only after re-sowing (significant year x land use interaction; fig. 3a). Surprisingly, experimental climate change had only a minor, marginally significant effect on ANPP, and neither the climate x land use interaction nor the three-way interaction with year were significant. However, when evaluating climate effects for each land-use type individually (table 1), we found that ANPP on EP was on average lower under future than under ambient climate, while ANPP on IM was lower under future climate in the first study years but higher in the second year of drought (fig. 3a).

**Fig. 3:**
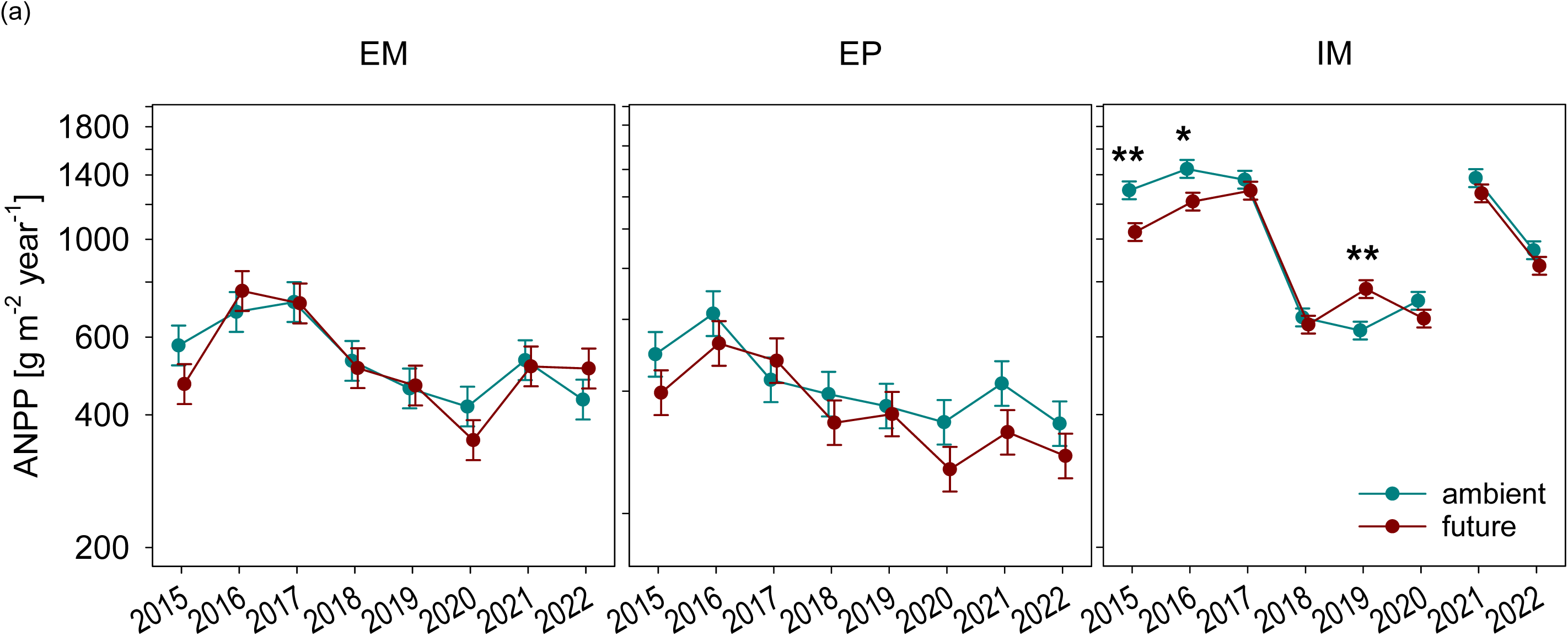
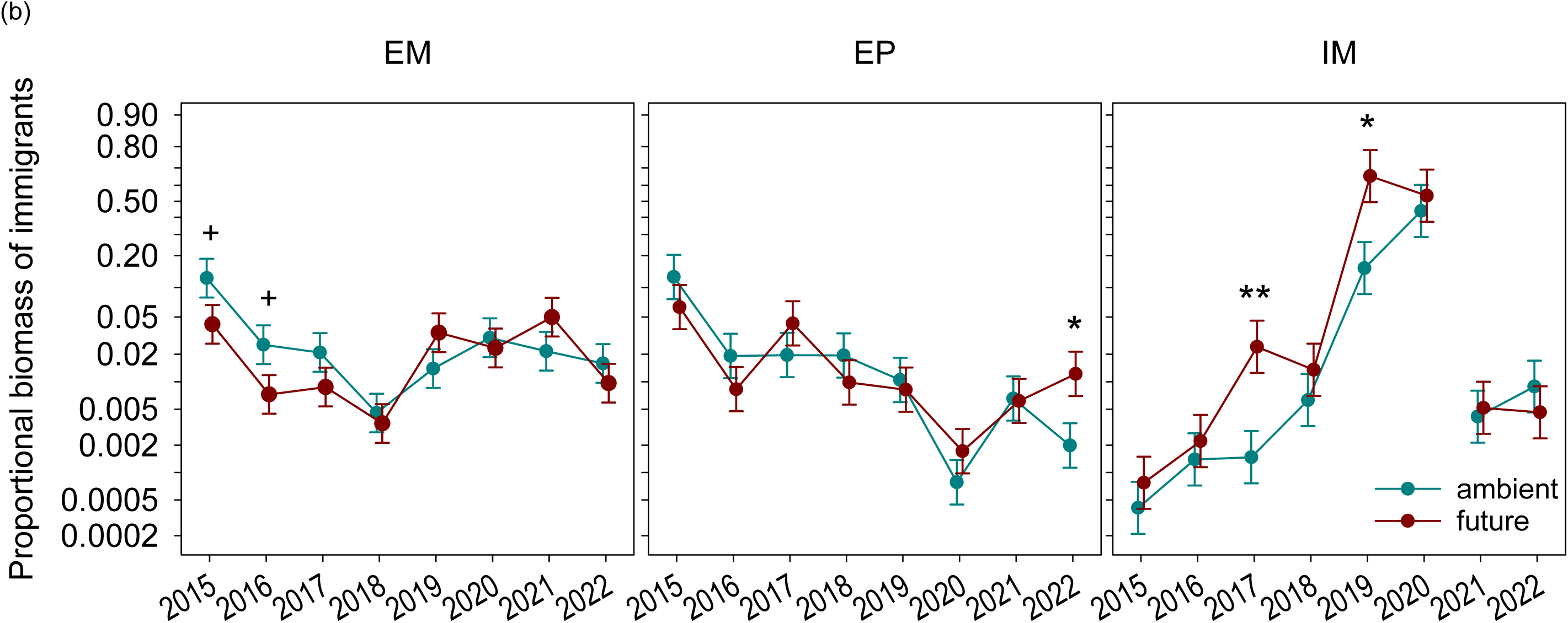
(a) Annual above-ground net primary production (ANPP, log scaled) and (b) proportional contribution of immigrant species to above-ground biomass (logit scaled) of extensively used meadows (EM), extensively used pastures (EP) and intensively used meadows (IM) under ambient and future climatic conditions across eight study years. Please note that IM was re-sown in fall 2020. (Marginal means and their standard errors extracted from generalized linear mixed effects models; significance of simple main effects of climate for each year and land-use type: ^+^ p < 0.1, * p < 0.05, ** p < 0.01, *** p < 0.001.)

The proportional contribution of immigrant species to aboveground biomass did, on average, not significantly differ between intensively and extensively used grasslands, but was larger on EM than on EP (table 1). However, proportional biomass of immigrants showed distinct temporal patterns according to land use (significant year x land use interaction; fig. 3b): On EM and EP plots, it decreased until the first and the third year of drought, respectively, and recovered afterwards. On IM plots, proportional biomass of immigrants continuously increased until the plots were re-sown. This was driven by a decreasing absolute biomass of residents during drought years and a continuously increasing immigrant biomass (fig. S3a, b; table S2). While there was no significant effect of climate manipulation on EM and EP, future climate led to an increase in the proportional biomass of immigrants on IM (significant climate x land use interaction: table 1; fig. 3b).

To evaluate whether the seasonal pattern of precipitation change affected response of ANPP to climate manipulation, we inspected intra-annual dynamics of standing biomass more closely (table 2, fig. S4). In all three grassland types, biomass reached its maximum in spring (first harvest on EM, first two harvests on EP and IM), during or just after experimentally increased precipitation under future climatic conditions. At that time, there was almost no difference in EM biomass between ambient and future climate, and a marginally significant decrease of EP biomass under future conditions. Future climate had a weak positive effect on IM biomass in early spring, but a significantly negative effect in late spring. In midsummer, after a period of reduced precipitation, biomass of EM and IM was lower under future than under ambient conditions, while EP showed no difference between climate treatments. Only IM biomass was harvested in fall, when it was still lower under future conditions despite increased precipitation.

**Table 2:** Results of repeated measurements ANOVAs based on generalized linear mixed effects models for the effects of climate, year and harvest date on seasonal above-ground biomass. Because of the different numbers of biomass harvests, separate models were compiled for each land use type (EM: extensively used meadows, EP: extensively used pastures, IM: intensively used meadows; d.f. enumerator and denominator degrees of freedom; F ratios: + p < 0.1, * p < 0.05, ** p < 0.01, *** p < 0.001).

|  | EM |  | EP |  | IM |  |
| --- | --- | --- | --- | --- | --- | --- |
|  | d.f. | F ratio | d.f. | F ratio | d.f. | F ratio |
| Climate | 1, 8 | 2.94 | 1, 8 | 1.65 | 1, 8 | 32.28 *** |
| Year | 7, 120 | 27.46 *** | 7, 154 | 9.33 *** | 7, 248 | 105.20 *** |
| Harvest | 1, 120 | 2053.58 *** | 2, 154 | 116.81 *** | 3, 248 | 559.70 *** |
| Year x Climate | 7, 120 | 0.35 | 7, 154 | 0.54 | 7, 248 | 4.37 *** |
| Harvest x Climate | 1, 120 | 22.48 *** | 2, 154 | 0.40 | 3, 248 | 14.24 *** |
| Year x Harvest | 7, 120 | 64.62 *** | 11, 154 | 9.54 *** | 19, 248 | 41.68 *** |
| Year x Harvest x Climate | 7, 120 | 1.71 | 11, 154 | 0.67 | 19, 248 | 1.45 |

### Effects of inter-annual climate variability on ANPP

In the two extensively used grasslands, relative difference in ANPP (as quantified by log response ratio, LRR) between the driest year 2018 and the wettest year 2021 was close to zero (EM: −0.002 ± 0.025, EP: −0.026 ± 0.074), and even smaller in magnitude than the LRR to experimental climate manipulation (EM: −0.021 ± 0.062, EP: −0.151 ± 0.046). ANPP on IM plots showed a much larger response to the difference in background climatic conditions (−0.316 ± 0.024) compared with its response to experimental climate manipulation (−0.067 ± 0.030). Furthermore, response to climate manipulation did not change with background climatic conditions of a particular year (main effect of dMI on LRR: F_1,12_ = 0.08, p = 0.78, interaction dMI x land use: F_2,12_ = 0.29, p = 0.75).

While ANPP varied independently of dMI on EM and EP, it increased strongly with increasing dMI, i.e. with decreasing drought, on IM (fig. 4, table S3). There was no main or interactive effect of climate manipulation on the relationship between ANPP and treatment-specific dMI, indicating that climate manipulation affected ANPP in the same way as background climate fluctuations.

**Fig. 4:**
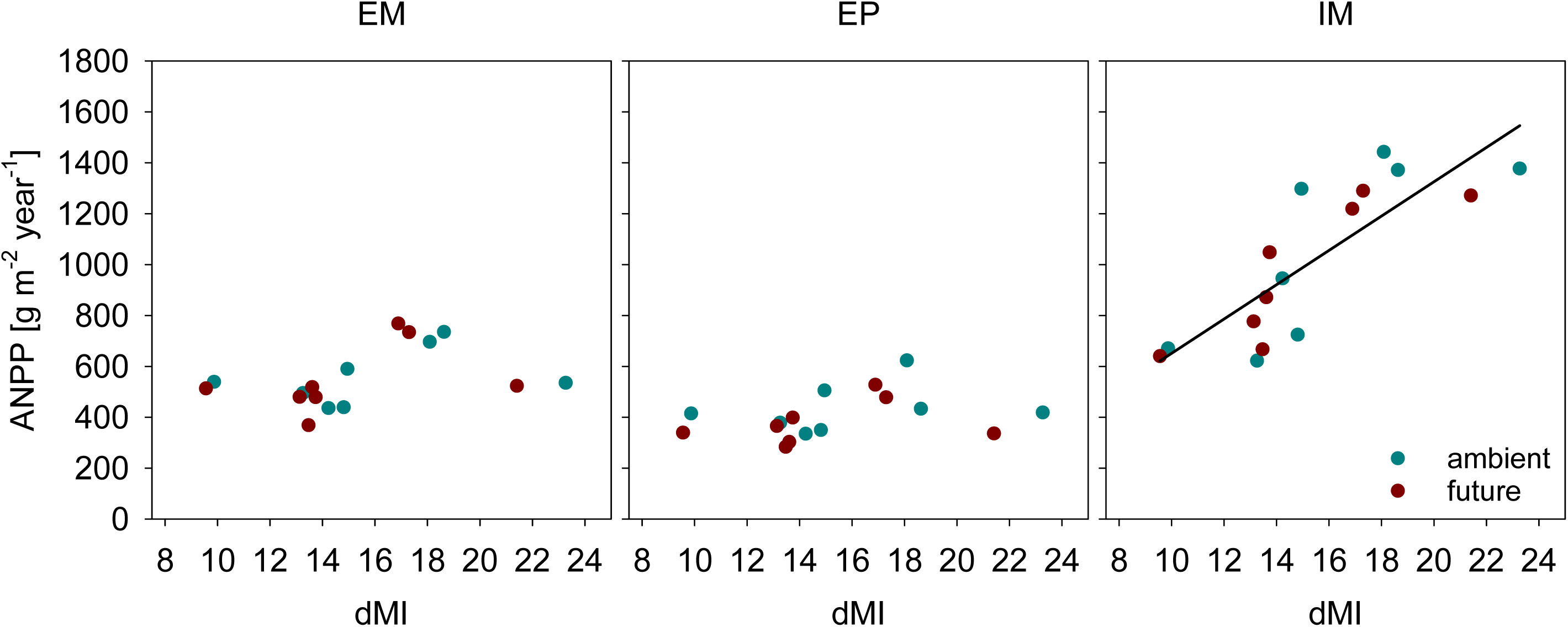
Effects of de Martonne’s index (dMI) and climate manipulation on annual above-ground net primary production (ANPP) of extensively used meadows (EM), extensively used pastures (EP) and intensively used meadows (IM) under ambient and future climatic conditions. The de Martonne’s index combines annual precipitation sum and mean annual temperature of each year and climate treatment, and increases with decreasing drought. According to ANCOVA, the climate x land use interaction is significant, but climate manipulation has no effect (see Appendix table 2). Separate regression analyses for each land-use type reveal that productivity does not significantly depend on dMI on EM (r^2^ = 0.16, p = 0.13) and EP (r^2^ = 0.11, p = 0.21), while the relationship is significant for IM (r^2^ = 0.67, p < 0.0001).

Although the resulting coefficient of variation of ANPP (as an inverse measure of stability) did not show a significant climate x land use interaction (table S4), a-priori contrasts revealed that the coefficient of variation was significantly higher (i.e. stability was lower) on IM compared to the two extensively used grasslands which did not differ from each other (fig. 5). However, this effect was only evident under ambient climate, while there was no difference among land use types under future climatic conditions.

**Fig. 5:**
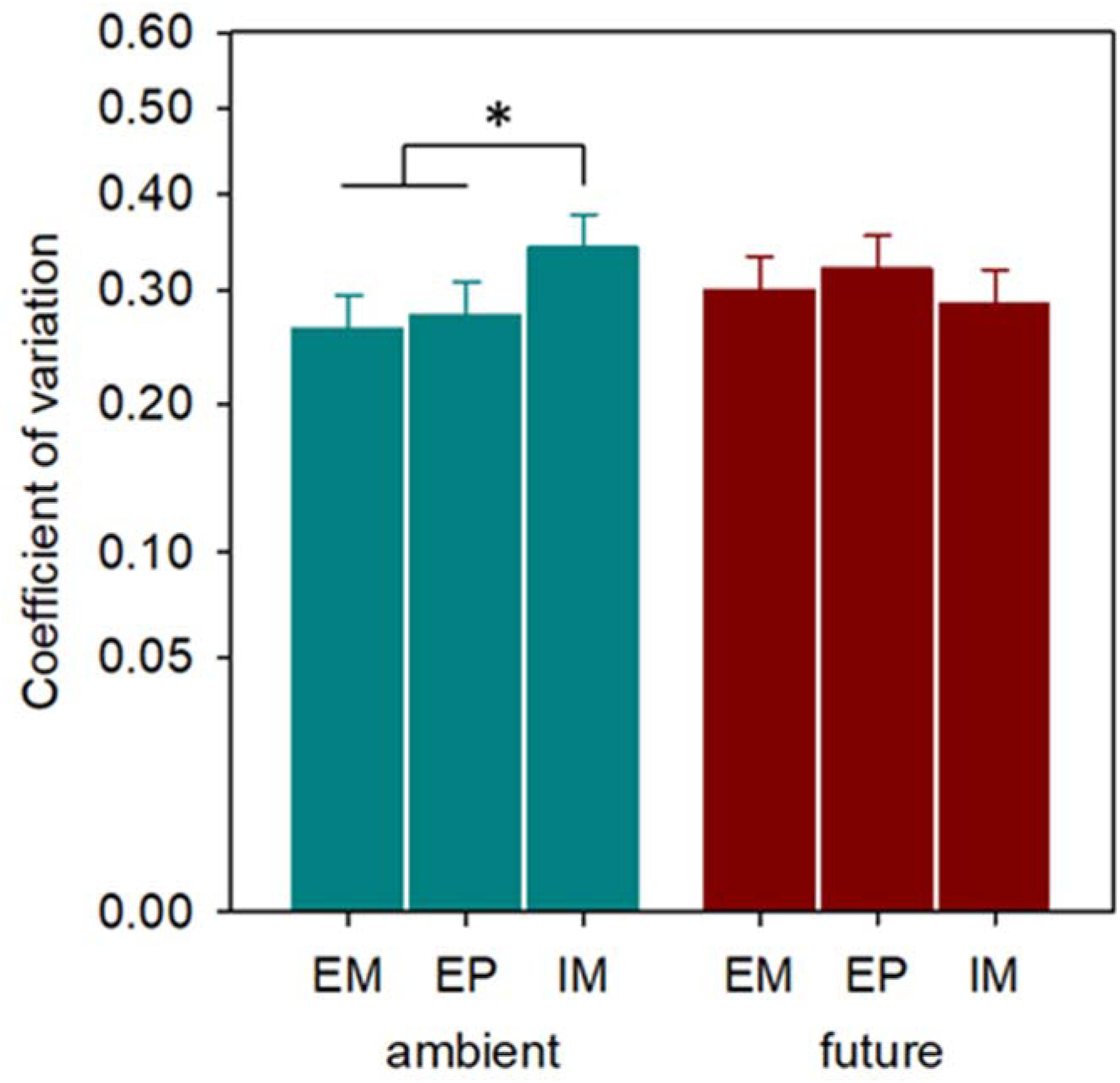
Coefficient of variation in annual above-ground net primary production (ANPP, square-root scaled) as an inverse measure of temporal stability of extensively used meadows (EM), extensively used pastures (EP) and intensively used meadows (IM) under ambient and future climatic conditions. (Marginal means and their standard errors extracted from generalized linear mixed effects models; contrast between the two extensively used grasslands and the intensively used grassland: * p < 0.05.)

## Discussion

While climate and land-use change are expected to interactively affect ecosystems, this interaction is rarely considered (Oliver and Morecroft 2014), especially in experimental studies. We found that the effects of inter-annual variability in background climate, including a series of extreme drought years, on the diversity and productivity of grasslands exceeded the effects of seasonal experimental climate manipulation that was superimposed onto background climate variability. Providing a high species diversity, extensively managed grasslands could better resist climate change compared to intensively used, species-poor grasslands.

The plant species diversity of the grasslands changed through time and weakly across experimental climate treatments. In line with our expectations, the diversity responses to climate manipulation and inter-annual climate variability were still stronger and more variable in intensively used grasslands. Initially, intensively used meadows were much less diverse than extensively used grasslands (meadows and pastures), as the former were composed of a few grass cultivars that are recommended for agricultural grasslands in drier regions in Central Germany (see Schädler et al. (2019) for more details). Changes in diversity were almost exclusively due to increases in the number of immigrant species, which may have entered from within (soil seed bank) or from outside (seed migration) the experiment. All grasslands showed higher richness and evenness as measured by the effective number of species in the future climate treatment in some years. This is likely because more dieback of resident species occurred in the future climate treatment, particularly in drought years, creating opportunities for establishment of drought-adapted immigrant species (see also Catford et al. 2020). Many other studies have also shown that less diverse communities are more likely to be invaded (e.g. Hector et al. 2001, Roscher et al. 2009). This pattern is often attributed to the higher number of unfilled niches compared to more diverse communities (MacArthur 1970, Levine and D’Antonio 1999). We would like to emphasize that the responses of our grasslands represent transitions of resident ecosystems in response to global change drivers, characterized by abundance shifts and immigration from the regional species pool (see Smith et al. 2009).

We found relatively weak effects of experimental climate on the aboveground net primary production (ANPP) of all grassland ecosystems. It should be noted that our climate manipulation retained the background inter-annual climate variability including the extremes. The weak overall response of productivity to the climate manipulation might be explained by the seasonal nature of the climate manipulation (see also Heitschmidt and Vermeire 2006, Denton et al. 2017, Hajek and Knapp 2022), in which spring and fall are wetter and summer is drier, according to the climate model projections. In our grasslands, maximum growth occurs in spring, and regrowth of plants during the summer contributes less to annual productivity. Thus, under future climate conditions the slight increases in productivity in spring due to higher water availability can compensate for the decline in productivity during the dry summer. Interestingly, the productivity of intensively used meadows in fall was still reduced under future climate even though this treatment involves higher fall precipitation, indicating a reduced capacity of these grasslands to recover from summer drought. There is a clear need for more experimental research on factors that potentially mediate post-drought ecosystem recovery (Knapp et al. 2023) such as land-use intensity.

At the beginning of the experiment, the ANPP of intensively used grasslands was high, and the future climate caused a small decrease in ANPP. As time passed, immigrants contributed increasingly to ANPP, especially in the future climate treatment and after the drought years, and overall productivity declined. Our results are in line with other studies that have found that fertile grasslands primarily composed of fast-growing species showed a stronger response to climate change compared to infertile grasslands (Grime et al. 2000, Van Sundert et al. 2021, but see Stampfli et al. 2018). The declining productivity and increase of immigrants in intensively used grasslands have practical implications for farmers, because these immigrant species have much lower fodder value compared to the originally sown cultivars (fodder values (mean ± SD) in 2020; sown: 8.67±0.33; immigrants: 3.60 ±0.52; https://wiki.ufz.de/biolflor/index.jsp, Klotz, Kühn, Durka 2002). The process of degradation, in which immigrant species replace the resident cultivars, is well known, and farmers expect to plow and re-sow their intensively used grasslands after some years (Bayerische Landesanstalt für Landwirtschaft 2018). However, we show here that with climate change, this process of degradation will be accelerated, which may impose additional economic costs as a consequence of a faster decreasing fodder value and the need for more frequent management measures.

During the course of the experiment, Central Europe experienced a series of consecutive droughts (years 2018, 2019 and 2020) that were amongst the most severe since the beginning of records (Boergens et al. 2020, Hari et al. 2020, Rakovec et al. 2022). We therefore had the unique opportunity to quantify the stability of differently used grasslands during a period with large inter-annual climate variability. We found a clear signal of the drought years on ANPP, especially in the intensively used grasslands for which we found a clear positive relationship between ANPP and water availability (quantified as dMI). Accordingly, intensively used grasslands had a larger coefficient of variation (CV) under ambient climate conditions, suggesting lower stability in ANPP compared to extensively used grasslands. As CV seems to decrease with increasing mean biomass in most cases (Döring et al. 2015), and in particular in grasslands (Reckling et al. 2021), the larger CV in our intensively managed, highly productive grassland can be considered a conservative estimate. Confirming our hypothesis, the results clearly show a lower resistance of intensively used grasslands to drought compared to the extensively used ones. Other studies showed that different aspects of land-use intensity, e.g. fertilization (Bharat et al. 2020, Meng et al. 2021, Van Sundert et al. 2021) or high mowing frequency (Vogel et al. 2012) decreased the resistance of grasslands to drought. Generally, these results are in line with the insurance hypothesis (Yachi and Loreau 1999) suggesting that diverse ecosystems have a greater capacity to adjust to fluctuations in resource availability compared to ecosystems with low diversity.

The stronger effects of inter-annual climate variability compared to experimental climate manipulation in our study might be due to the intensity of the drought events, which were unprecedented in their intensity (Rakovec et al. 2022) and likely exceeded a threshold of tolerance in water scarcity for most plants. Two meta-analyses also report stronger responses of plant communities to inter-annual climate variability compared to climate manipulation (Langley et al. 2018, Kröel-Dulay et al. 2022). Our seasonal climate manipulation was intended to correspond to a realistic future climate scenario, and was not as extreme as many other experimental climate studies (Korell et al. 2020). Nevertheless, even when the magnitudes of experimental manipulation are stronger weak responses of plant productivity to climate manipulations are quite common (Grime et al. 2000, Jentsch et al. 2007, Jentsch et al. 2011, Hoover et al. 2014). When considering belowground processes, other published studies from the GCEF have demonstrated that future climate conditions reduced soil biological activity (Siebert et al. 2019, Kostin et al. 2021) and changed soil microbial communities (Bei et al. 2023). These alterations in belowground processes can feedback on aboveground processes such as productivity (Bardgett & Van der Putten 2014) and emphasize the need to jointly consider above- and below-ground processes in global change experiments in order to understand related ecosystem functions.

## Conclusions

Climate and land-use change are known to be major threats for biodiversity and functioning of ecosystems, but how these global change drivers interact is less clear. Our study provides experimental evidence that land-use intensity alters the response of grassland communities to climate change. A realistic scenario of climate change, simulating seasonal shifts of precipitation, had relatively small effects on diversity and productivity of grasslands which were most pronounced in intensively used species-poor grasslands. Future climate accelerated the shift from valuable forage cultivars to less valuable plant communities composed primarily of immigrating species, which will have economic consequences for farmers. Likewise, intensively used grasslands showed significantly lower resistance to inter-annual climate variability, including the most severe droughts since the beginning of records, even though seed mixtures included cultivars recommended for dry regions. We therefore conclude that extensive management, together with other measures to increase species diversity of grasslands, is an important tool to stabilize primary production under the impact of climate change.

## Supporting information

Land-use intensity mediates response to climate_SuppInfo

## Acknowledgements

We appreciate the Helmholtz Association, the Federal Ministry of Education and Research, the State Ministry of Science and Economy of Saxony-Anhalt and the State Ministry for Higher Education, Research and the Arts Saxony to fund the Global Change Experimental Facility (GCEF) project. We acknowledge François Buscot and Thomas Reitz for their role in setting up the GCEF. We thank Thomas Bienert, Sven Meyer, Eckhard Winter and especially Konrad Kirsch for their work in in maintaining the plots and infrastructures of the Global Change Experimental Facility (GCEF). We thank Alexandra Weigelt for their contributions to the experiment, as well as Antje Thondorf, Birgit Müller, Birgit Sawall, Charlotte Beber, Christa Wolfram, Christine Voigt, Franz Dussl, Ina Geier, Katrin Kittlaus, Martina Herrmann, Renate Hintz, Roman Patzak, Sabine Straßenburg, Steffi Wagner, Verena Schmidt and Wladimir Gelfgat as well as numerous students for their help with harvesting or sorting biomass samples. Furthermore, we thank Guido Warthemann and Thomas Premper (LPR Landschaftsplanung Dr. Reichhoff GmbH) as well as Jens Stolle for contributing to the vegetation censuses.

## Author contributions

L. Korell: Conceptualization; formal analysis, investigation, writing – original draft and preparation, writing – review and editing; H. Auge: conceptualization, data curation, formal analysis, funding acquisition, investigation, methodology, writing – original draft and preparation, writing – review and editing; M. Schädler: methodology, project administration (scientific coordination), writing – review and editing; I. Merbach: project administration (technical coordination), investigation, writing – review and editing; S. Klotz: funding acquisition, methodology, writing – review and editing; S. Berger: data curation, investigation, writing – review and editing; W. Durka, S. Haider, C. Roscher, E. Welk: methodology, investigation, writing – review and editing; M. Andrzejak, I. Hensen, Y. Herion, J. Höfner, L. Kindermann, T.M. Knight, A. Linstädter, A-M. Madaj, S. Michalski and C. Plos investigation, writing – review and editing.

## Notes

### Competing Interest Statement

The authors have declared no competing interest.

## References

Alexander, J. M., J. M. Diez, and J. M. Levine. 2015. Novel competitors shape species’ responses to climate change. Nature 525:515–518.

Altermann, M., J. Rinklebe, I. Merbach, et al. 2005. Chernozem—Soil of the Year 2005. Journal of Plant Nutrition and Soil Science 168:725–740.

Andrade, C., J. Contente, and J. A. Santos. 2021. Climate change projections of aridity conditions in the Iberian Peninsula. Water 13:2035.

Bai, Y., and M. F. Cotrufo. 2022. Grassland soil carbon sequestration: Current understanding, challenges, and solutions. Science 377:603–608.

Bardgett, R. D., & W. H. Van Der Putten. 2014. Belowground biodiversity and ecosystem functioning. Nature 515:505–51.

Bastazini, V. A., N. Galiana, H. Hillebrand, et al. 2021. The impact of climate warming on species diversity across scales: Lessons from experimental meta- ecosystems. Global Ecology and Biogeography 30:1545–1554.

Bayerische Landesanstalt für Landwirtschaft. 2018. Saatguteinsatz im Grünland. Übersaat - Nachsaat – Neuansaat. Landesanstalt für Landwirtschaft und Gartenbau Sachsen-Anhalt.

Beckmann, M., K. Gerstner, M. Akin-Fajiye, et al. 2019. Conventional land-use intensification reduces species richness and increases production: A global meta-analysis. Global Change Biology 25:1941–1956.

Bellard, C., C. Bertelsmeier, P. Leadley, et al. 2012. Impacts of climate change on the future of biodiversity. Ecology Letters 15:365–377.

Bengtsson, J., J. M. Bullock, B. Egoh, et al. 2019. Grasslands—more important for ecosystem services than you might think. Ecosphere 10:e02582.

Bharath, S., E. T. Borer, L. A. Biederman, et al. 2020. Nutrient addition increases grassland sensitivity to droughts. Ecology 101:e02981.

Biurrun, I., R. Pielech, I. Dembicz, et al. 2021. Benchmarking plant diversity of Palaearctic grasslands and other open habitats. Journal of Vegetation Science 32:e13050.

Boergens, E., A. Güntner, H. Dobslaw, et al. 2020. Quantifying the Central European Droughts in 2018 and 2019 With GRACE Follow-On. Geophysical Research Letters 47:e2020GL087285.

Catford, J. A., J. M. Dwyer, E. Palma, et al. 2020. Community diversity outweighs effect of warming on plant colonization. Global Change Biology 26:3079–3090.

Chase, J. M., B. J. McGill, D. J. McGlinn, et al. 2018. Embracing scale-dependence to achieve a deeper understanding of biodiversity and its change across communities. Ecology Letters 21:1737–1751.

Compagnoni, A., S. Levin, D. Z. Childs, et al. 2021. Herbaceous perennial plants with short generation time have stronger responses to climate anomalies than those with longer generation time. Nature Communications 12:1–8.

Davy, R., Esau, I., A. Chernokulsky et al. 2017 Diurnal asymmetry to the observed global warming. International Journal of Climatology 37:79–93.

Denton, E. M., J. D. Dietrich, M. D. Smith, et al. 2017. Drought timing differentially affects above-and belowground productivity in a mesic grassland. Plant Ecology 218:317–328.

Döring, T. F., S. Knapp, and J. E. Cohen. 2015. Taylor’s power law and the stability of crop yields. Field Crops Research 183:294–302.

Döring, T. F., and M. Reckling. 2018. Detecting global trends of cereal yield stability by adjusting the coefficient of variation. European Journal of Agronomy 99:30–36.

Fay, P. A., J. D. Carlisle, A. K. Knapp, et al. 2003. Productivity responses to altered rainfall patterns in a C4-dominated grassland. Oecologia 137:245–251.

Gherardi, L. A., and O. E. Sala. 2019. Effect of interannual precipitation variability on dryland productivity: A global synthesis. Global Change Biology 25:269–276.

Gibson, D. J., and J. A. Newman. 2019. Grasslands and climate change. Cambridge University Press.

Gornish, E. S., and J. M. Tylianakis. 2013. Community shifts under climate change: mechanisms at multiple scales. American Journal of Botany 100:1422–1434.

Gossner, M. M., T. M. Lewinsohn, T. Kahl, et al. 2016. Land-use intensification causes multitrophic homogenization of grassland communities. Nature 540:266–269.

Grime, J. P., V. K. Brown, K. Thompson, et al. 2000. The response of two contrasting limestone grasslands to simulated climate change. Science 289:762–765.

Gruner, D. S., M. E. Bracken, S. A. Berger, et al. 2017. Effects of experimental warming on biodiversity depend on ecosystem type and local species composition. Oikos 126:8–17.

Hajek, O. L., and A. K. Knapp. 2022. Shifting seasonal patterns of water availability: ecosystem responses to an unappreciated dimension of climate change. New Phytologist 233:119–125.

Hari, V., O. Rakovec, Y. Markonis, et al. 2020. Increased future occurrences of the exceptional 2018–2019 Central European drought under global warming. Scientific Reports 10:1–10.

Hastings, A. 2004. Transients: the key to long-term ecological understanding? Trends in Ecology & Evolution 19:39–45.

Harvey, L. D. D. 1995. Warm days, hot nights. Nature 377:15–16

Hector, A., K. Dobson, A. Minns, et al. 2001. Community diversity and invasion resistance: an experimental test in a grassland ecosystem and a review of comparable studies. Ecological Research 16:819–831.

Hedges, L. V., J. Gurevitch, and P. S. Curtis. 1999. The meta-analysis of response ratios in experimental ecology. Ecology 80:1150–1156.

Heitschmidt, R., and L. Vermeire. 2006. Can abundant summer precipitation counter losses in herbage production caused by spring drought? Rangeland Ecology & Management 59:392–399.

Hill, M. O. 1973. Diversity and evenness: a unifying notation and its consequences. Ecology 54:427–432.

Hoover, D. L., A. K. Knapp, and M. D. Smith. 2014. Resistance and resilience of a grassland ecosystem to climate extremes. Ecology 95:2646–2656.

Hurlbert, S. H. 1971. The nonconcept of species diversity: a critique and alternative parameters. Ecology 52:577–586.

IPCC. 2021. Climate Change 2021: The Physical Science Basis. Contribution of Working Group I to the Sixth Assessment Report of the Intergovernmental Panel on Climate Change. Cambridge University Press, Cambridge, United Kingdom and New York, NY, USA.

Isbell, F., D. Craven, J. Connolly, et al. 2015. Biodiversity increases the resistance of ecosystem productivity to climate extremes. Nature 526:574–577.

Jentsch, A., J. Kreyling, and C. Beierkuhnlein. 2007. A new generation of climate- change experiments: events, not trends. Frontiers in Ecology and the Environment 5:365–374.

Jentsch, A., J. Kreyling, M. Elmer, et al. 2011. Climate extremes initiate ecosystem-regulating functions while maintaining productivity. Journal of Ecology 99:689–702.

Jost, L. 2006. Entropy and diversity. Oikos 113: 363–375.

Kirmer, A., and S. Tischew. 2006. Handbuch naturnahe Begrünung von Rohböden. Springer.

Klotz, S., Kühn, I., and W. Durka ed. 2002. BIOLFLOR: Eine Datenbank mit biologisch-ökologischen Merkmalen zur Flora von Deutschland. Schriftenreihe für Vegetationskunde 38:1–334. Bundesamt für Naturschutz, Bonn.

Knapp, A. K., M. L. Avolio, C. Beier, et al. 2017. Pushing precipitation to the extremes in distributed experiments: recommendations for simulating wet and dry years. Global Change Biology 23:1774–1782.

Knapp, A. K., A. Chen, R. J. Griffin-Nolan, et al. 2020. Resolving the Dust Bowl paradox of grassland responses to extreme drought. Proceedings of the National Academy of Sciences 117:22249–22255.

Knapp, A. K., Condon, K. V., C. C Folks et al. 2023. Field experiments have enhanced our understanding of drought impacts on terrestrial ecosystems— But where do we go from here?. Functional Ecology.

Komatsu, K. J., M. L. Avolio, N. P. Lemoine, et al. 2019. Global change effects on plant communities are magnified by time and the number of global change factors imposed. Proceedings of the National Academy of Sciences 116:17867–17873.

Korell, L., H. Auge, J. M. Chase, et al. 2020. We need more realistic climate change experiments for understanding ecosystems of the future. Global Change Biology 26:325–327.

Korell, L., H. Auge, J. M. Chase, et al. 2021. Responses of plant diversity to precipitation change are strongest at local spatial scales and in drylands. Nature Communications 12:1–7.

Kostin, J. E., S. Cesarz, A. Lochner, et al. 2021. Land-use drives the temporal stability and magnitude of soil microbial functions and modulates climate effects. Ecological Applications 31:e02325.

Kreyling, J., Arfin Khan, M. A., F. Sultana, et al. 2017. Drought effects in climate change manipulation experiments: quantifying the influence of ambient weather conditions and rain-out shelter artifacts. Ecosystems 20: 301–315.

Kröel-Dulay, G., A. Mojzes, K. Szitár, et al. 2022. Field experiments underestimate aboveground biomass response to drought. Nature Ecology & Evolution 6:540–545.

Landesamt für Landwirtschaft, U. u. l. R. 2010. Beweidung von Offen-und Halboffenbiotopen. Eine adäquate Pflegemethode unter besonderer Berücksichtigung der FFH-Lebensraumtypen und Arten. Schriftenreihe des Landesamtes für Landwirtschaft, Umwelt und ländliche Räume des Landes Schleswig-Holstein 18:1–30.

Landesanstalt für Landwirtschaft und Gartenbau Sachsen-Anhalt. 2020. Mischungs-und Sortenempfehlungen Grünland, Bernburg, Germany.

Langley, J. A., S. K. Chapman, K. J. La Pierre, et al. 2018. Ambient changes exceed treatment effects on plant species abundance in global change experiments. Global Change Biology 24:5668–5679.

Levine, J. M., and C. M. D’Antonio. 1999. Elton revisited: a review of evidence linking diversity and invasibility. Oikos 87:15–26.

MacArthur, R. H. 1970. Theoretical Population Biology 1:1–11.

McNaughton, S.J, D. G. Milchunas, and D. A. Frank. 1996. How can net primary productivity be measured in grazing ecosystems? Ecology 77:974–977.

Meng, B., J. Li, G. E. Maurer, et al. 2021. Nitrogen addition amplifies the nonlinear drought response of grassland productivity to extended growing-season droughts. Ecology 102:e03483.

Morris, W. F., J. Ehrlén, J. P. Dahlgren, et al. 2020. Biotic and anthropogenic forces rival climatic/abiotic factors in determining global plant population growth and fitness. Proceedings of the National Academy of Sciences 117:1107–1112.

Naumann, G., Cammalleri, C., L. Mentaschi et. al. 2021. Increased economic drought impacts in Europe with anthropogenic warming. Nature Climate Change 11:485–491.

O’Mara, F. P. 2012. The role of grasslands in food security and climate change. Annals of Botany 110:1263–1270.

Oliver, T. H., and M. D. Morecroft. 2014. Interactions between climate change and land use change on biodiversity: attribution problems, risks, and opportunities. WIREs Climate Change 5:317–335.

Parmesan, C. 2006. Ecological and evolutionary responses to recent climate change. Annual Review of Ecology, Evolution, and Systematics:637–669.

Piao, S., Q. Liu, A. Chen, et al. 2019. Plant phenology and global climate change: Current progresses and challenges. Global Change Biology 25:1922–1940.

Pimm, S. L. 1984. The complexity and stability of ecosystems. Nature 307:321–326.

Poschlod, P., J. P. Bakker, and S. Kahmen. 2005. Changing land use and its impact on biodiversity. Basic and Applied Ecology 6:93–98.

Poschlod, P., A. Baumann, and P. Karlik. 2009. Origin and development of grasslands in Central Europe. Pages 15–25, Grasslands in Europe. KNNV Publishing.

Rakovec, O., L. Samaniego, V. Hari, et al. 2022. The 2018–2020 Multi-year drought sets a new benchmark in Europe. Earth’s Future 10:e2021EF002394.

Regionaler-Klimaatlas (https://www.regionaler-klimaatlas.de/), online 19.07.2023

Reckling, M., H. Ahrends, T.-W. Chen, et al. 2021. Methods of yield stability analysis in long-term field experiments. A review. Agronomy for Sustainable Development 41:1–28.

Renner, S. S., and C. M. Zohner. 2018. Climate change and phenological mismatch in trophic interactions among plants, insects, and vertebrates. Annual Review of Ecology, Evolution, and Systematics 49:165–182.

Roscher, C., A. J. Fergus, J. S. Petermann, et al. 2013. What happens to the sown species if a biodiversity experiment is not weeded? Basic and Applied Ecology 14:187–198.

Roscher, C., V. M. Temperton, N. Buchmann, et al. 2009. Community assembly and biomass production in regularly and never weeded experimental grasslands. Acta Oecologica 35:206–217.

Ruppert, J. C., and A. Linstädter. 2014. Convergence between ANPP estimation methods in grasslands—A practical solution to the comparability dilemma. Ecological Indicators 36:524–531.

Sabatini, F. M., B. Jiménez-Alfaro, U. Jandt, et al. 2022. Global patterns of vascular plant alpha diversity. Nature Communications 13:1–16.

Sala, O. E., F. Stuart Chapin, J. J. Armesto, et al. 2000. Global biodiversity scenarios for the year 2100. Science 287:1770–1774.

Sandel, B., L. J. Goldstein, N. J. Kraft, et al. 2010. Contrasting trait responses in plant communities to experimental and geographic variation in precipitation. New Phytologist 188:565–575.

Schädler, M., F. Buscot, S. Klotz, et al. 2019. Investigating the consequences of climate change under different land-use regimes: a novel experimental infrastructure. Ecosphere 10:e02635.

Schils, R. L., C. Bufe, C. M. Rhymer, et al. 2022. Permanent grasslands in Europe: Land use change and intensification decrease their multifunctionality. Agriculture, Ecosystems & Environment 330:107891.

Siebert, J., M. P. Thakur, T. Reitz, et al. 2019. Extensive grassland-use sustains high levels of soil biological activity, but does not alleviate detrimental climate change effects. Advances in Ecological Research 60: 25–58.

Smith, M. D., A. K. Knapp, and S. L. Collins. 2009. A framework for assessing ecosystem dynamics in response to chronic resource alterations induced by global change. Ecology 90:3279–3289.

Song, J., S. Wan, S. Piao, et al. 2019. A meta-analysis of 1,119 manipulative experiments on terrestrial carbon-cycling responses to global change. Nature Ecology & Evolution 3:1309–1320.

Speißer, B., Wilschut, R. A., and Van Kleunen, M. 2022. Number of simultaneously acting global change factors affects composition, diversity and productivity of grassland plant communities. Nature Communications 13: 7811.

Stampfli, A., J. M. Bloor, M. Fischer, et al. 2018. High land-use intensity exacerbates shifts in grassland vegetation composition after severe experimental drought. Global Change Biology 24:2021–2034.

Tälle, M., B. Deák, P. Poschlod, et al. 2016. Grazing vs. mowing: A meta-analysis of biodiversity benefits for grassland management. Agriculture, Ecosystems & Environment 222:200–212.

Vogel, A., M. Scherer-Lorenzen, and A. Weigelt. 2012. Grassland resistance and resilience after drought depends on management intensity and species richness. PLOS ONE 7:e36992.

van Sundert, K., Arfin Khan, M. A., S. Bharath et al. (2021). Fertilized graminoids intensify negative drought effects on grassland productivity. Global Change Biology 27: 2441–2457.

von Ende, C. N. 2001. Repeated-measures analysis. In Design and Analysis of Ecological Experiments (pp. 134–157). Oxford University Press.

Warton, D. I., and F. K. Hui. 2011. The arcsine is asinine: the analysis of proportions in ecology. Ecology 92:3–10.

White, R. P., S. Murray, M. Rohweder, et al. 2000. Grassland ecosystems. World Resources Institute Washington, DC, USA.

Wilcox, K. R., J. C. von Fischer, J. M. Muscha, et al. 2015. Contrasting above- and belowground sensitivity of three Great Plains grasslands to altered rainfall regimes. Global Change Biology 21:335–344.

Woodward, J. A., and D. G. Bonett. 1991. Simple main effects in factorial designs. Journal of Applied Statistics 18:255–264.

Wu, Z., P. Dijkstra, G. W. Koch, et al. 2011. Responses of terrestrial ecosystems to temperature and precipitation change: A meta-analysis of experimental manipulation. Global Change Biology 17:927–942.

Yachi, S., and M. Loreau. 1999. Biodiversity and ecosystem productivity in a fluctuating environment: the insurance hypothesis. Proceedings of the National Academy of Sciences 96:1463–1468.

