## Supplementary material for "Low land-use intensity buffers grasslands against future climate and inter-annual climate variability in a large field experiment": Land-use intensity mediates response to climate_SuppInfo

### Supporting information A: Figures and tables

**Table S1:** List of the 56 plant species that were sown in early spring 2014 to establish the extensively used meadows (EM) and extensively used pastures (EP) in the Global Change Experimntal Facility (GCEF)

1. Grasses

*Anthoxanthum odoratum* L.
*Arrhenatherum elatius* (L.) P. Beauv. ex J. Presl & C. Presl
*Brachypodium pinnatum* (L.) P. Beauv.
*Bromus erectus* Huds.
*Bromus hordeaceus* L.
*Cynosurus cristatus* L.
*Dactylis glomerata* L.
*Festuca pratensis* Huds.
*Festuca rubra* L.
*Festuca rupicola* Heuff.
*Helictotrichon pratense* (L.) Besser
*Phleum pratens*e L.
*Poa angustifolia* L.
*Trisetum flavescens* (L.) P. Beauv.

2. Legumes

*Lathyrus pratensis* L.
*Lotus corniculatus* L.
*Medicago falcata* L.
*Medicago lupulina* L.
*Securigera varia* (L.) Lassen
*Trifolium campestre* Schreb.
*Trifolium pratense* L.
*Trifolium repens* L.
*Vicia cracca* L.
*Vicia sepium* L.

3. Non-legume forbs

*Achillea millefolium* L.
*Agrimonia eupatoria* L.
*Allium vineale* L.
*Campanula rapunculoides* L.
*Centaurea jacea* L.
*Crepis biennis* L.
*Daucus carota* L.
*Dianthus carthusianorum* L.
*Falcaria vulgaris* Bernh.
*Filipendula vulgaris* Moench
*Galium album* Mill.
*Galium verum* L.
*Geranium pratense* L.
*Heracleum sphondylium* L.
*Hypochaeris radicata* L.
*Knautia arvensis* (L.) Coult.
*Leontodon autumnalis* L.

*Leucanthemum vulgare* Lam. / *L. ircutianum* DC.
*Linaria vulgaris* Mill.
*Lychnis flos-cuculi* L.
*Pastinaca sativa* L.
*Pimpinella saxifraga* L.
*Plantago lanceolata* L.
*Plantago media* L.
*Prunella vulgaris* L.
*Ranunculus acris* L.
*Salvia pratensis* L.
*Scabiosa ochroleuca* L.
*Silene latifolia* Poir.
*Stachys recta* L.
*Tragopogon orientalis* L.
*Veronica chamaedrys* L.

**Table S2:** Results of repeated measurements ANOVAs based on generalized linear mixed effects models for the effects of climate, land use and year on richness of resident species, proportional richness of immigrant species, effective number of species $S_{PIE}$, above-ground biomass of resident species and above-ground biomass of immigrant species. To test a-priori hypotheses, the main effect of land use is decomposed into orthogonal contrasts, and the climate x land use interaction is decomposed into simple main effects of climate on each land-use type (EM: extensively used meadows, EP: extensively used pastures, IM: intensively used meadows; d.f. enumerator and denominator degrees of freedom; + p < 0.1, * p < 0.05, ** p < 0.01, *** p < 0.001).

| | | F ratio | | | | |
|---|---|---|---|---|---|---|
| | d.f. | Richness of residents | Proportional richness of immigrants | $S_{PIE}$ | Biomass of residents | Biomass of immigrants |
| Climate | 1, 8 | 0.24 | 2.29 | 2.21 | 2.01 | 0.25 |
| Land use | 2, 16 | 20700.70 *** | 587.12 *** | 2265.94 *** | 13.88 *** | 6.77 ** |
| Extensive vs. Intensive | 1, 16 | 39501.80 *** | 1147.24 *** | 4355.47 *** | 5.32 * | 0.80 |
| EM vs. EP | 1, 16 | 54.09 *** | 1.88 | 1.99 | 22.43 *** | 12.74 ** |
| Climate x Land use | 2, 16 | 2.55 | 0.07 | 6.52 ** | 5.31 * | 3.55 + |
| Climate \| EM | 1,16 | 2.69 | 0.84 | 9.21 ** | 0.04 | 0.83 |
| Climate \| EP | 1,16 | 0.22 | 0.80 | 0.13 | 0.18 | 0.04 |
| Climate \| IM | 1,16 | 0.83 | 1.01 | 0.46 | 8.73 ** | 3.60 + |
| Year | 7, 56 | 66.11 *** | 41.95 *** | 43.92 *** | 43.77 *** | 5.18 *** |
| Year x Climate | 7, 56 | 1.37 | 2.10 + | 1.64 | 0.87 | 1.55 |
| Year x Land use | 14, 112 | 12.75 *** | 18.46 *** | 139.32 *** | 14.37 *** | 21.74 *** |
| Year x Climate x Land use | 14, 112 | 0.80 | 1.53 | 1.16 | 1.34 | 1.50 |

**Table S3:** Results of ANCOVA based on generalized linear mixed effects model on effects of de Martonne's index (dMI), climate manipulation, and land use on annual above-ground net primary production (ANPP) averaged across replicate plots (d.f. enumerator and denominator degrees of freedom; F ratios: + p < 0.1, * p < 0.05, ** p < 0.01, *** p < 0.001).

| | d.f. | F ratio |
|---|---|---|
| dMI | 1, 29 | 6.80 * |
| Climate | 1, 29 | 0.15 |
| Land use | 2, 29 | 4.73 * |
| Climate x Land use | 2, 29 | 0.12 |
| dMI x Climate | 1, 29 | 0.05 |
| dMI x Land use | 2, 29 | 32.53 *** |
| dMI x Climate x Land-use | 2, 29 | 0.14 |

**Table S4:** Results of ANOVA based on generalized linear mixed effects model on the effects of climate and land use on the coefficient of variation of net primary production (ANPP). To test a-priori hypotheses, the climate x land use interaction is decomposed into contrasts between land-use types at each climate treatment (d.f. enumerator and denominator degrees of freedom; F ratios: + p < 0.1, * p < 0.05, ** p < 0.01, *** p < 0.001).

| | d.f. | F ratio |
|---|---|---|
| Climate | 1, 8 | 0.07 |
| Land use | 2, 16 | 0.82 |
| Climate x Land use | 2, 16 | 2.35 |
| extensive vs. intensive \| ambient | 1, 16 | 5.33 * |
| extensive meadow vs. extensive pasture \| ambient | 1, 16 | 0.13 |
| extensive vs. intensive \| future | 1, 16 | 0.57 |
| extensive meadow vs. extensive pasture \| future | 1, 16 | 0.32 |

**Fig. S1:** Annual precipitation sum and mean annual temperature of the experimental site during the study period. (Dotted line: long-term mean 1896-2013.)

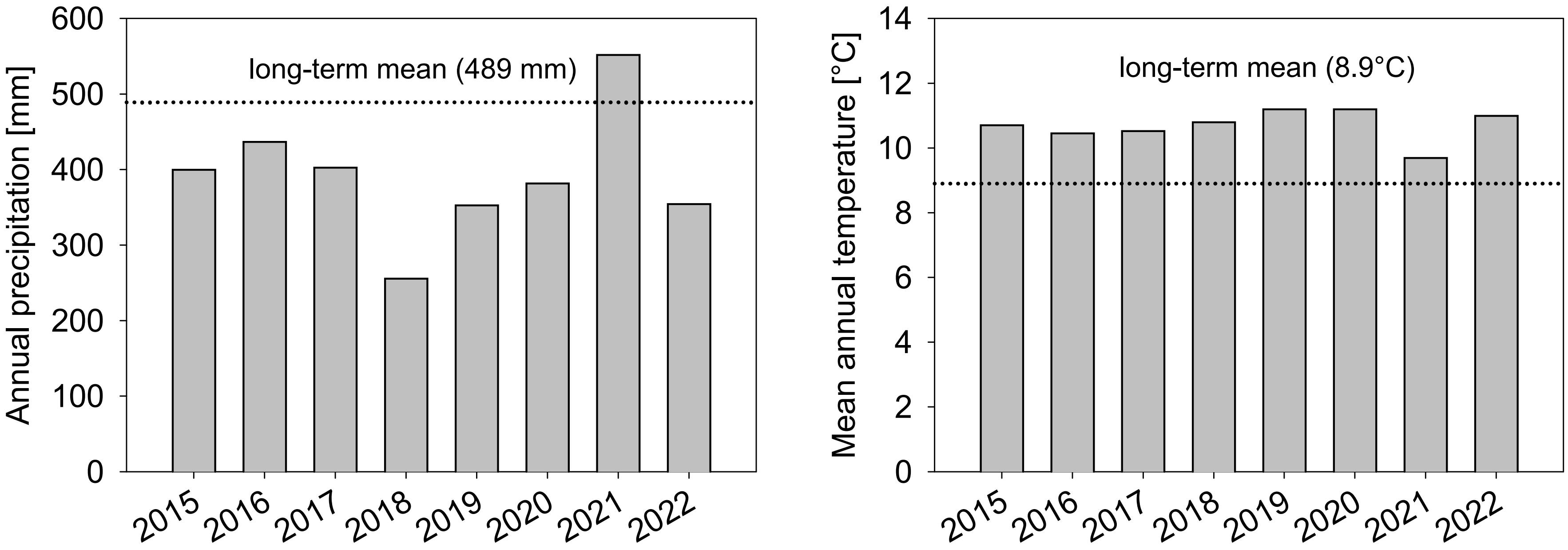

**Fig. S2:** a) Richness of resident species (log scaled), (b) proportional richness of immigrant species (logit scaled), and (c) effective number of species $S_{PIE}$ (log scaled) of extensively used meadows (EM), extensively used pastures (EP) and intensively used meadows (IM) under ambient and future climatic conditions across eight study years, as measured on 9 $m^2$ sampling plots. Please note that IM was re-sown in fall 2020. (Marginal means and their standard errors extracted from generalized linear mixed effects models; significance of simple main effects of climate for each year and land-use type: + $p < 0.1$, * $p < 0.05$, ** $p < 0.01$, *** $p < 0.001$.)

(a)

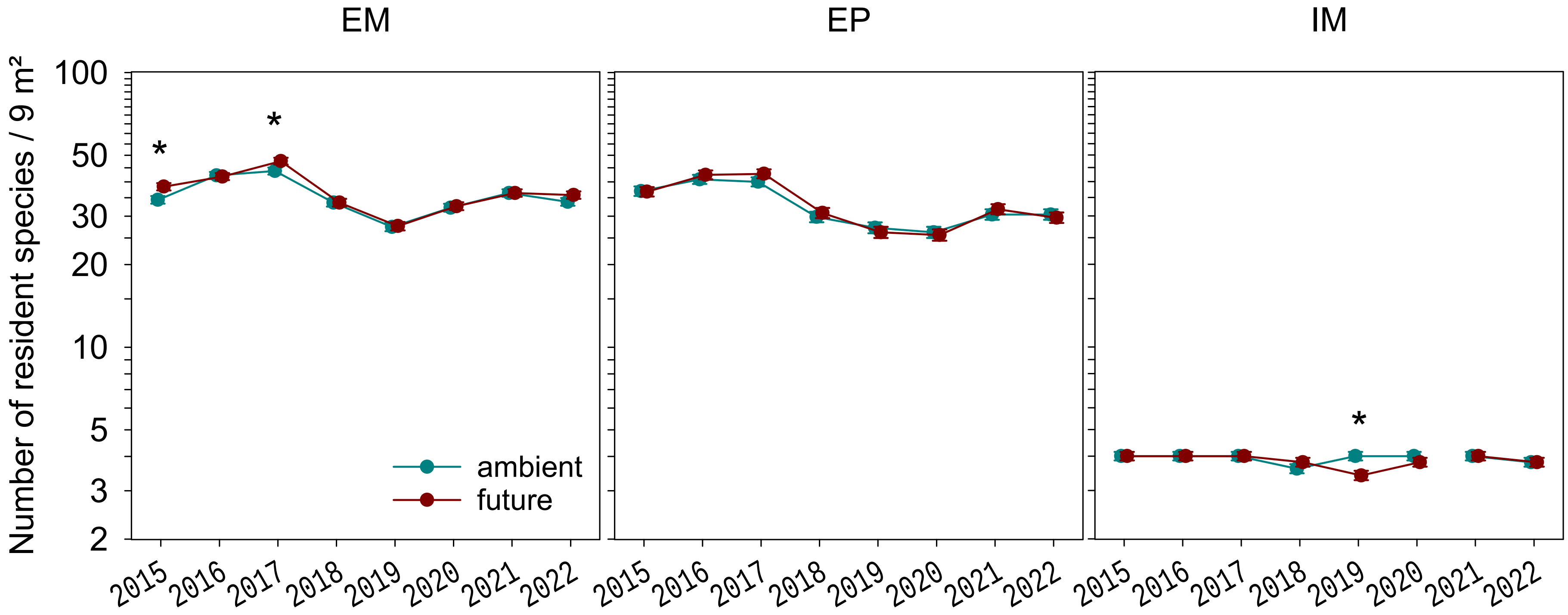

(b)

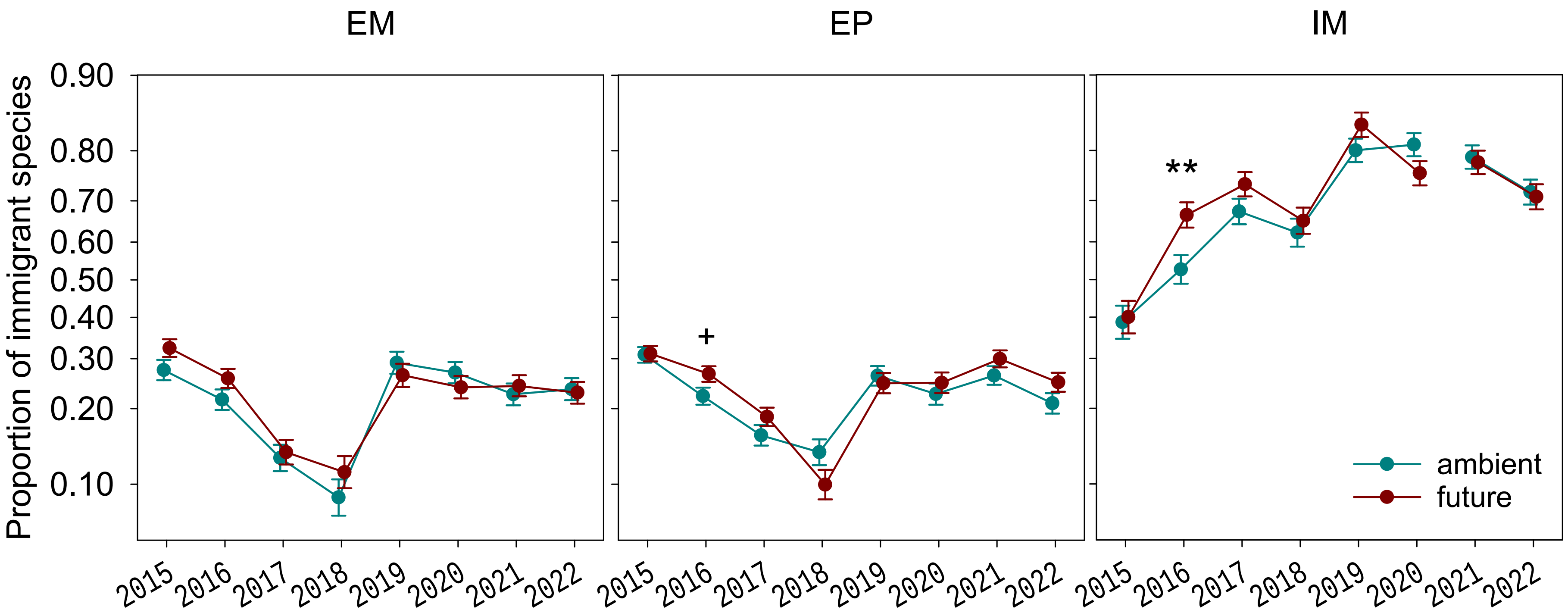
EM
EP
IM
Proportion of immigrant species
0.90
0.80
0.70
0.60
0.50
0.40
0.30
0.20
0.10
+
**
ambient
future
2015
2016
2017
2018
2019
2020
2021
2022

(c)

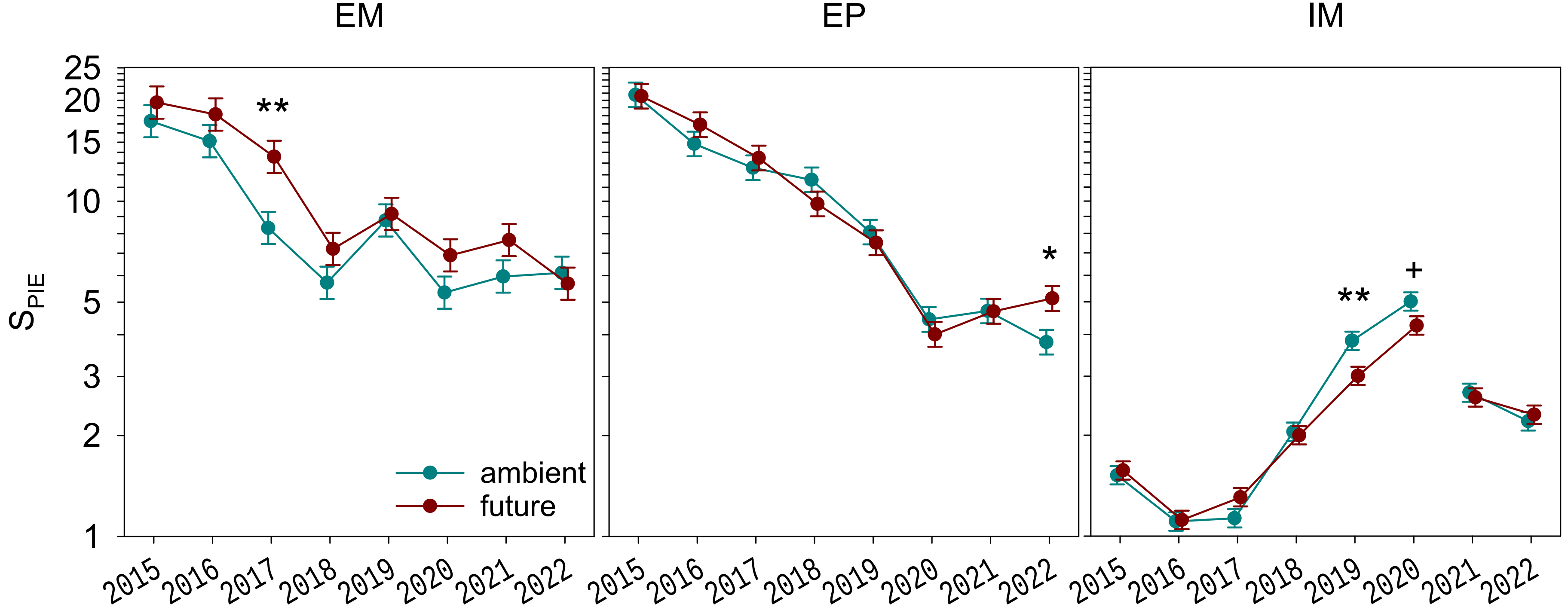

EM
EP
IM
$S_{PIE}$
25
20
15
10
5
3
2
1
**
*
**
+
ambient
future
2015
2016
2017
2018
2019
2020
2021
2022

**Fig. S3:** (a) Above-ground biomass of resident species and (b) above-ground biomass of immigrant species of extensively used meadows (EM), extensively used pastures (EP) and intensively used meadows (IM) under ambient and future climatic conditions across eight study years (y axes log scaled). Please note that IM was re-sown in fall 2020. (Marginal means and their standard errors extracted from generalized linear mixed effects models; significance of simple main effects of climate for each year and land-use type: + p < 0.1, * p < 0.05, ** p < 0.01, *** p < 0.001.)

(a)

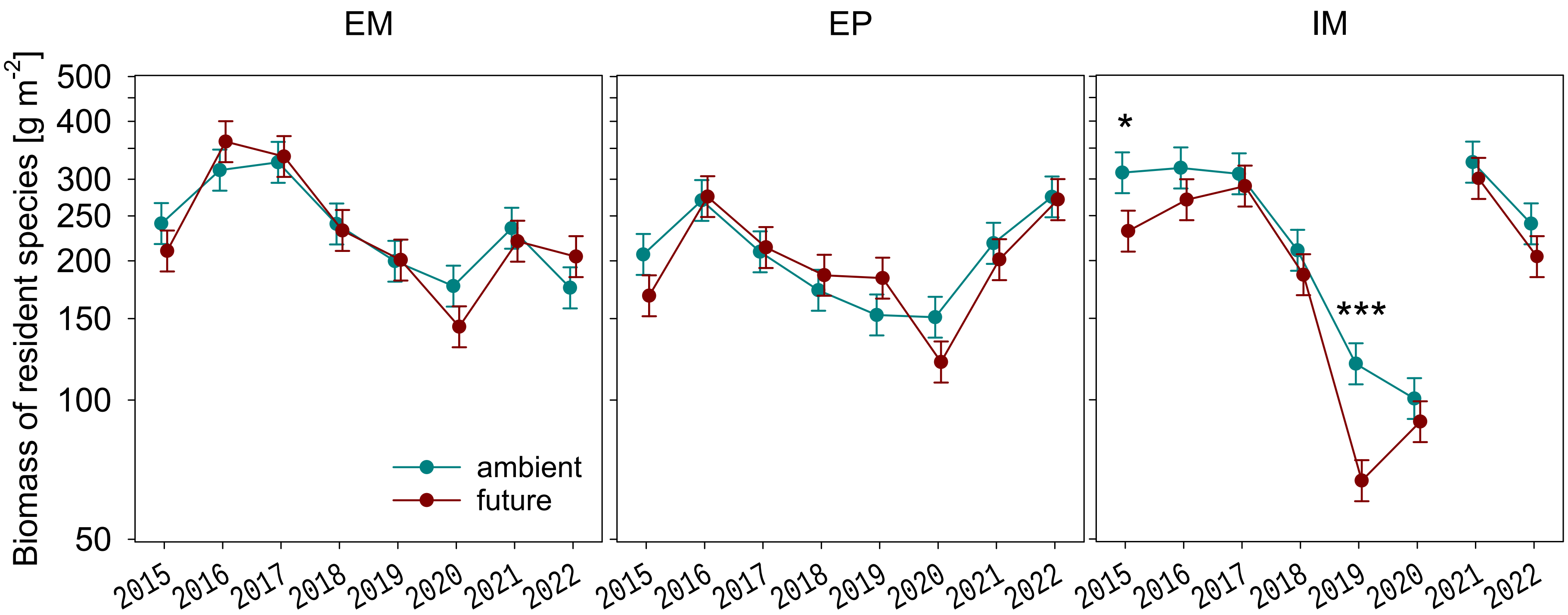

(b)

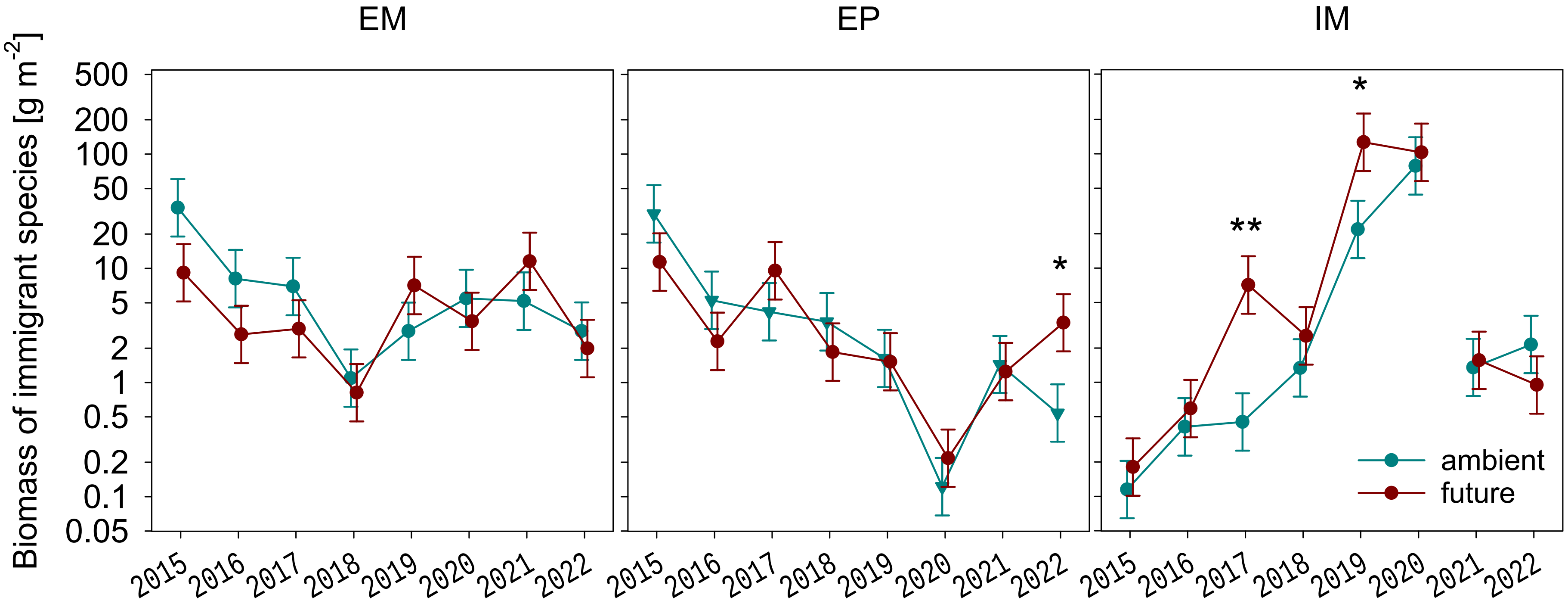
EM
EP
IM
Biomass of immigrant species [g m-2]
500
200
100
50
20
10
5
2
1
0.5
0.2
0.1
0.05
2015
2016
2017
2018
2019
2020
2021
2022
*
**
*
ambient
future

**Fig. S4:** Above-ground standing biomass (log scaled) of each harvest date on extensively used meadows (EM), extensively used pastures (EP) and intensively used meadows (IM) under ambient and future climatic conditions, averaged across eight study years. Please note that standing biomass on the extensively used pastures does only partially reflect regrowth, as on average 66% of aboveground biomass remain after each grazing (range 42% – 84% across harvest dates and years). (Marginal means and their standard errors extracted from generalized linear mixed effects models; significance of simple main effects of climate for each harvest date and land-use type: + $p < 0.1$, * $p < 0.05$, ** $p < 0.01$, *** $p < 0.001$.)

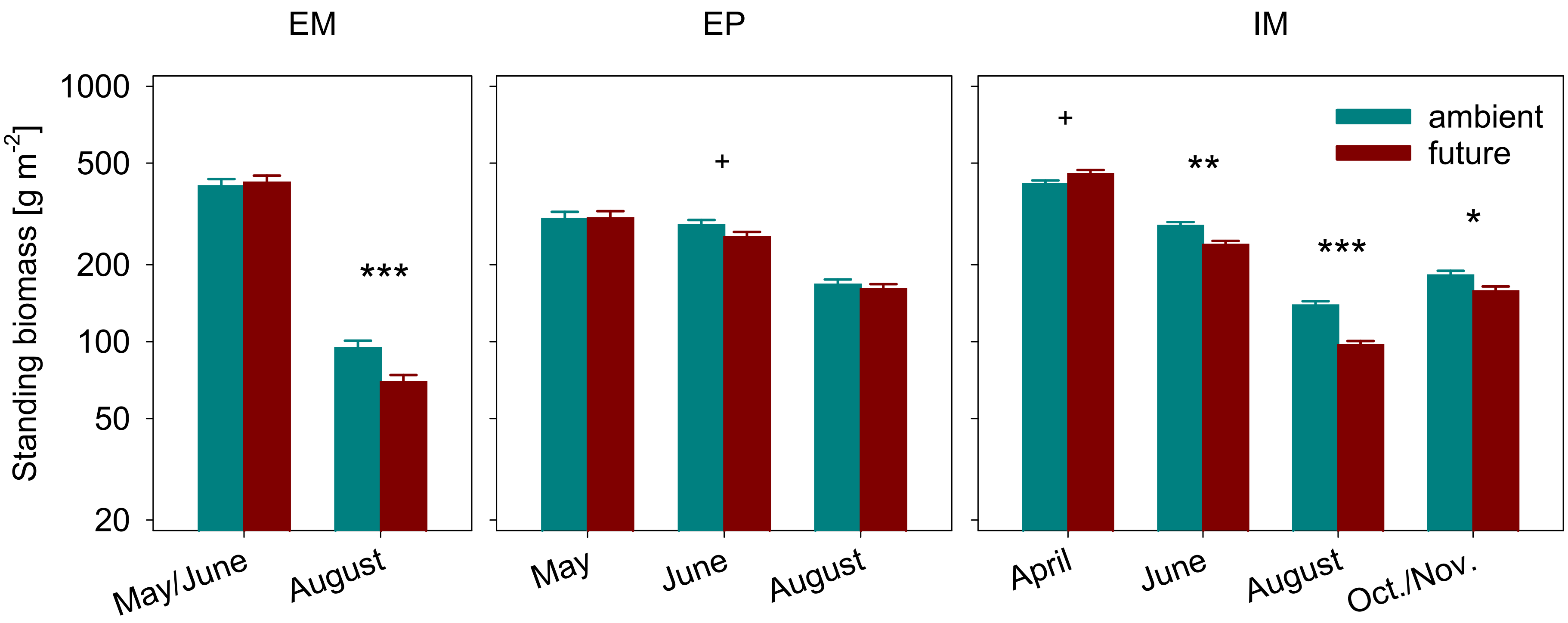
